# PRSS56 acts as an intrinsic retinal signal driving postnatal ocular axial growth and myopia susceptibility

**DOI:** 10.64898/2026.01.11.697334

**Authors:** Kiran Gangappa, Durairaj Duraikannu, Jayshree Advani, Cassandre Labelle-Dumais, Emre Berk Uludag, Nivedita Singh, Swanand Koli, Chen Jiang, Yien-Ming Kuo, Yoshihiro Ishikawa, Hélène Choquet, Anand Swaroop, K. Saidas Nair

## Abstract

Myopia is a leading cause of visual impairment worldwide, and high myopia markedly increases the risk of irreversible vision loss. Although visual experience guides postnatal ocular elongation, the role of intrinsic retinal growth signals remains poorly defined. Here we identify the serine protease PRSS56 as a retinal factor that promotes ocular axial growth beyond early development. Using genetic mouse models, we show that conditional inactivation of Prss56 in Müller glia reduces axial length and causes hyperopia even under dark-rearing conditions, demonstrating that PRSS56 drives axial elongation independently of light-evoked visual input during emmetropization. Conversely, Müller glia-specific overexpression of Prss56 induces axial elongation in a proteolysis-dependent manner, supporting its role as an autonomous retinal growth signal. In concordance, human genetic analyses reveal that the common *PRSS56* variant rs2853447 is associated with increased axial length in individuals with myopia and high myopia, but not in non-myopes, suggesting that this variant confers a selective growth advantage in individuals predisposed to ocular elongation. Functional genomic analyses further identify a myopia-associated variant, rs2741297, within a retinal enhancer in intron 4 of *PRSS56* marked by open chromatin and transcription factor occupancy. Together, these findings establish PRSS56 as an intrinsic retinal growth factor that functions beyond early eye development and support a model in which genetic and environmental factors converge on retinal pathways to modulate myopia susceptibility.

## Introduction

Myopia is now a prevalent refractive disorder worldwide, and its incidence is rising rapidly.^1,2^ Yet, effective interventions to prevent or slow down myopia progression currently remain limited.^3,4^ Corrective lenses or refractive surgery can help restore clear vision; however, these measures do not address the underlying causes of abnormal ocular axial growth or mitigate the risk of associated complications.^5,6^ Notably, myopia is a major risk factor for blinding diseases such as retinal detachment, myopic maculopathy, and glaucoma and is projected to become a leading cause of blindness by 2050.^7,8^ This underscores an urgent need to identify therapeutic strategies to modulate axial elongation and restore healthy refractive development thereby preventing vision impairment associated with myopia pathogenesis.^4,9^ Its rampant rise is largely attributed to lifestyle changes, including reduced time spent outdoors.^10^ Nonetheless, myopia has a strong genetic basis and is highly heritable. Elucidating how the genetic predisposition and environmental influences interact to drive myopia remains a central challenge in the field.^10^

The precise regulation of ocular growth is critical to achieve normal vision, and ocular axial length constitutes a central determinant of emmetropia, the state where light focuses directly on the retina, resulting in clear vision. Notably, insufficient or excessive ocular axial elongation are well established causes of refractive errors, namely hyperopia and myopia, respectively. Ocular axial growth is governed through a cascade of signaling events, originating from the retina and relayed to the sclera. Retinal signaling induces scleral extracellular matrix (ECM) remodeling to promote ocular axial elongation^11–14^ and can be broadly divided into two categories: vision-unadjusted and vision-adjusted.^15^ Vision-unadjusted ocular growth is an intrinsic genetically programmed process that operates independently of the visual input during prenatal and postnatal developmental stages.^15^ On the other hand, emmetropization is a vision-guided postnatal developmental process that modulates ocular growth to fine-tune eye’s axial length to match its optical power.^10,16,17^ Additionally, ocular axial growth is known to be influenced by the spectral composition of light, with specific wavelengths and patterns of light reported to either promote or suppress axial elongation.^18,19^

The human eye undergoes a rapid growth phase that is largely driven by intrinsic growth mechanisms during early childhood. The eye typically grows from approximately 15 to 16 millimeters (mm) at birth to about 22 mm, reaching roughly >90 % of its adult size, by three years of age. Over the next 15 years or so, the eye continues to grow at a slower pace, leading to a modest 1-2 mm additional increase in ocular size.^20–23^ Although vision-guided emmetropization begins during infancy, its role becomes predominant during the later slow phase when ocular axial growth is fine-tuned by visual experience, enabling precise alignment of the optical focal plane to achieve emmetropia. The emmetropization process is well conserved across species, including in mice.^24^ However, the molecular and cellular mechanisms that regulate ocular axial growth remain poorly understood. Moreover, it is unknown whether intrinsic growth programs operate only during early development or also contribute to later stages, when ocular growth is predominantly dictated by visual experience. The mechanisms underlying prenatal and early rapid phase of ocular growth during infancy are presumed to be distinct from those regulating vision-guided postnatal growth. To date, no molecular factor regulating early ocular growth has been shown to remain active during later stages or to influence myopia susceptibility.

Genetic studies in humans have provided critical insights into the regulation of ocular growth, identifying multiple loci associated with refractive errors.^25–27^ Among these, *PRSS56*, a trypsin-like serine protease, has emerged as a particularly compelling candidate. Notably, common noncoding *PRSS56* variants have been strongly associated with increased ocular axial length and myopia in the general population.^25,26,28,29^ Conversely, autosomal recessive *PRSS56* mutations have been identified as a major cause of nanophthalmos, a condition characterized by severely shortened axial length and extreme hyperopia.^30–34^ In concordance, we previously showed that the loss of *PRSS56* function induces shorter ocular axial length in mice.^30^ In a conditional knockout mouse model, *PRSS56* inactivation at distinct time points (ranging from postnatal day (P) 6 to P18) before the completion of retinal maturation results in reduced ocular axial length.^33^ The requirement of PRSS56 before the onset of visual experience (i.e. before P13, which corresponds to eye opening in mice) supports its role as an intrinsic driver of ocular growth.^33^ Collectively, these evidences position PRSS56 as a critical mediator of ocular axial growth and suggested that altered PRSS56 expression might contribute to refractive errors. In this scenario, reduced or elevated PRSS56 levels/activity could lead to insufficient or excessive ocular axial elongation, causing hyperopia or myopia, respectively. Interestingly, *Prss56* expression is upregulated by optical defocus paradigms (minus lens) which induce ocular elongation and myopia in marmosets, suggesting that *PRSS56* might also contribute to vision-guided ocular axial elongation postnatally in the mature retina as well.^35^ However, it remains unclear whether *PRSS56* can causally induce ocular elongation at a timeframe once the retina has matured and visual input becomes the dominant regulatory signal. Additionally, how common *PRSS56* variants contribute to myopia remains poorly defined.

In this study, we leveraged both mouse and human genetics to demonstrate that PRSS56 promotes ocular axial elongation independently of light-driven visual input during later developmental stages when the retina is fully mature and responsive to visual cues. We demonstrate that selective overexpression of PRSS56 in mouse Müller glia is sufficient to drive ocular axial elongation. We have also uncovered genetic association of a common *PRSS56* variant previously linked to ocular axial length with increased axial length in myopic, but not emmetropic, individuals. Finally, integration of genetic association and epigenomic data has revealed that *PRSS56* risk variants are located within putative regulatory element(s) of *PRSS56* in the retina, indicating a possible mechanism of myopia susceptibility by non-coding genetic variations modulating *PRSS56* expression. Our findings establish retinal PRSS56 as a critical component of the ocular growth machinery and implicate its dysregulation in excessive axial length elongation associated with myopia. Thus, PRSS56 may represent a potential therapeutic target to slow excessive ocular axial growth and reduce the risk of vision loss in myopia.

## Results

### Müller glia-derived PRSS56 promotes ocular axial elongation during postnatal developmental stages when the retina is fully developed

We previously demonstrated that PRSS56 contributes to ocular axial growth during early postnatal eye development in the mouse.^30,33^ To determine whether PRSS56 contributes to ocular axial growth once the retina is fully mature and visual experience becomes the predominant regulator of ocular axial length, we utilized conditional Prss56 mutant mice. First, we crossed mice carrying a floxed *Prss56* allele (*Prss56^fl/fl^*) to the tamoxifen-inducible ubiquitous *Ubc-Cre^ERT2^* transgenic line for conditional inactivation of *Prss56* at 1 month (mo) of age, when the retina is functionally mature and emmetropization is actively taking place in the mouse.^16,36–38^ We performed ocular biometry at multiple timepoints before and after tamoxifen induction. Before tamoxifen administration at 1 mo, all ocular parameters examined, including axial length, vitreous chamber depth (VCD), lens diameter, and retinal thickness, were comparable between *Prss56^fl/fl^;Ubc^Cre^*and control *Prss56^fl/fl^* mice (**Fig.S1, Table S1**). However, following *Prss56* inactivation at 1 mo, we observed a modest but statistically significant decreases in axial length and VCD, along with a concurrent increase in retinal thickness in *Prss56^fl/fl^;Ubc^Cre^* mice aged to 2 mo and 4–5 mo (**Fig. S1**). These results demonstrate a continuous requirement for PRSS56 for ocular growth, not only during early stages of development but even at later time points when the retina is fully matured and axial growth is influenced by visual experience.

Since Müller glia-derived PRSS56 participates in promoting ocular axial growth^33^, we next selectively inactivated *Prss56* in Müller glial cells at 1 mo using the inducible *Glast*-*Cre^ERT^* line. Prior to tamoxifen injection, all ocular biometric parameters examined were comparable between *Prss56^fl/fl^;Glast^Cre^* mice and control littermates (**Fig. 1A-G and Table S2**). However, following tamoxifen induction at 1 mo, *Prss56^fl/fl^;Glast^Cre^* mice exhibited significant shortening in axial length and VCD along with an increase in retinal thickness compared to *Prss56^fl/fl^* littermates or tamoxifen uninjected *Prss56^fl/fl^;Glast^Cre^* control mice at both ages examined (2 mo and 3 mo). Consistent with these changes, tamoxifen-injected *Prss56^fl/fl^;Glast^Cre^* mice also exhibited hyperopic refractive shift and a robust increase in retinal *Adamts19* expression (a secreted metalloproteinase that remodels the ECM ), a molecular marker associated with PRSS56 loss-of-function^34^ that further confirm functional inactivation of the PRSS56-dependent pathway (**Fig. 1H–I**). To assess whether conditional inactivation of PRSS56 caused any structural or pathological abnormalities, we performed fundoscopy, fluorescein angiography and histological analyses. The retinal vasculature of tamoxifen-injected *Prss56^fl/fl^;Glast^Cre^* mice remained intact and appeared normal, and no gross morphological ocular or retinal defects were observed (**Fig. S2**). These findings validate that Müller glia-derived PRSS56 is responsible for continuous modulation of ocular growth that extends beyond early developmental windows.

**Figure 1.**
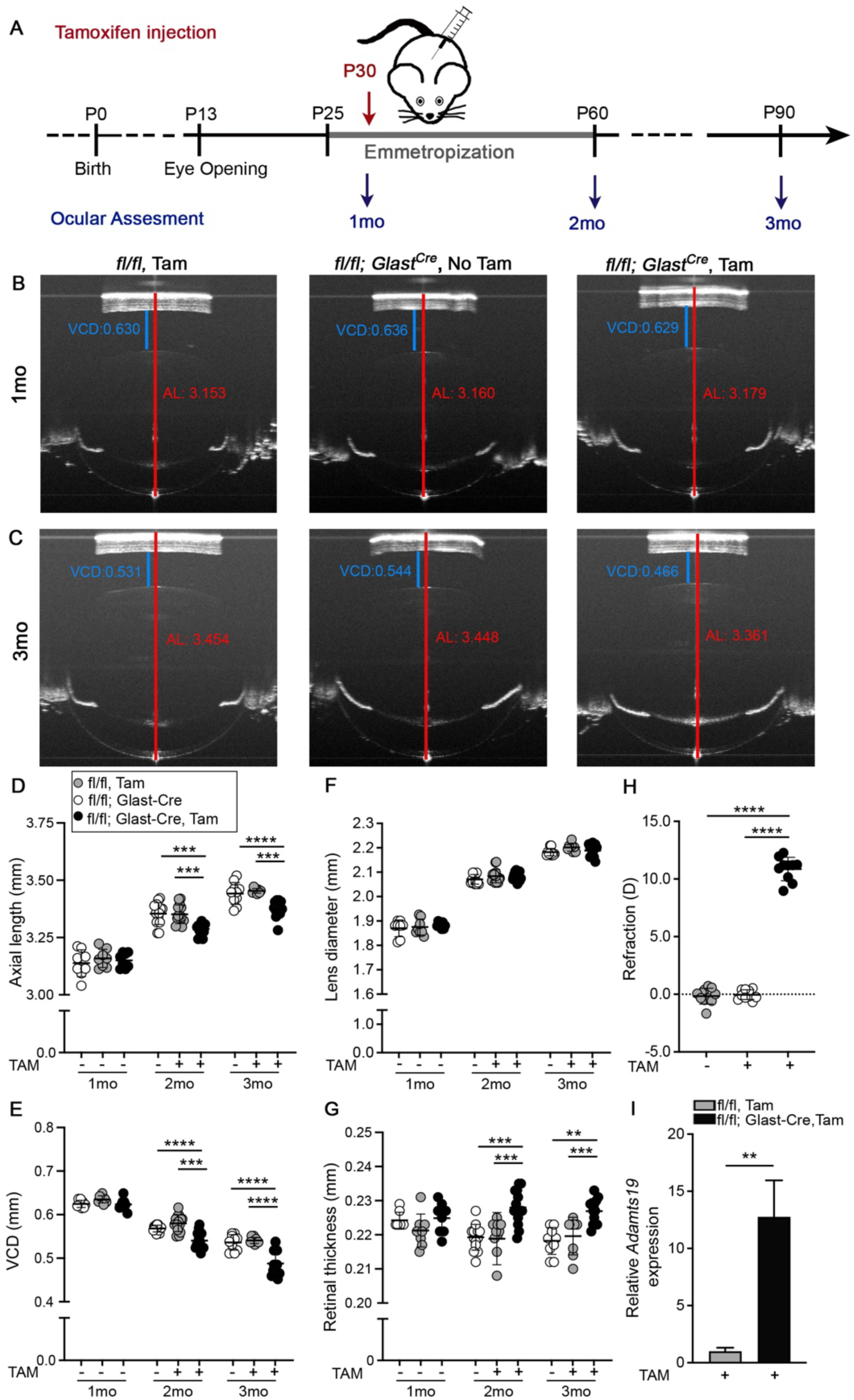
Selective inactivation of *Prss56* in Müller glia at 1 month of age leads to a shortened ocular axial length. *Prss56* was selectively ablated in Müller glia at 1 month of age (mo) by crossing *Prss56^fl/fl^* mice to the inducible *Glast-Cre^ERT^* mouse strain (*Glast-Cre*). Conditional Cre expression in Müller glial cells was induced by tamoxifen injection (TAM) at 1mo. (A-C) Representative OCT images from 1mo, 2mo and 3mo mice showing reduced axial length and vitreous chamber depth (VCD) and increased retinal thickness following conditional inactivation of *Prss56* in Müller glia at 1 mo (compare eyes from tamoxifen injected *Prss56^fl/fl^; Glast^Cre^* mice in B and C to those from injected *Prss56^fl/fl^* littermates or uninjected *Prss56^fl/fl^; Glast^Cre^* control mice). (D-G) Ocular biometric parameters were similar between *Prss56^fl/fl^* and *Prss56^fl/fl^;Glast^Cre^* mice prior to tamoxifen injection at 1mo. Following tamoxifen injection, significant reductions in ocular axial length (D) and VCD (E) and increase in retinal thickness (G) were detected at in *Prss56^fl/fl^;Glast^Cre^* mice compared to *Prss56^fl/fl^* littermates or uninjected control *Prss56^fl/fl^;Glast^Cre^* mice at both ages examined (2mo and 3mo). n= 8 eyes/group at 1mo; n= 14 eyes/group at 2mo; n= 14 injected and 10 uninjected *Prss56^fl/fl^; Glast^Cre^* eyes and 8 injected control *Prss56^fl/fl^* eyes at 3mo. (H) Consistent with reduced ocular axial length, *Prss56^fl/fl^; Glast^Cre^* mice injected with tamoxifen at 1mo display a hyperopic refraction compared to uninjected *Prss56^fl/fl^; Glast^Cre^* mice or injected control *Prss56^fl/fl^* littermates at 3 mo. n= 10 eyes/group. (I) *Adamts19* expression was also significantly increased in retina from 3mo *Prss56^fl/fl^; Glast^Cre^*mice compared to *Prss56^fl/fl^* littermates. n= 3 *Prss56^fl/fl^* and 4 *Prss56^fl/fl^;Glast^Cre^* eyes. Data are presented as mean ± SD; ** p<0.01; *** p<0.001; **** p<0.0001, One-Way ANOVA. Additional ocular parameter measurements are presented in Supplemental Table 2.

To further define the contribution of PRSS56 toward the end of the emmetropization period, which is typically completed around 2 mo in the mouse^24^, we next performed selective inactivation of *Prss56* from Müller glia at 2 mo. As expected, ocular parameters were similar between *Prss56^fl/fl^;Glast^Cre^* and *Prss56^fl/fl^* control mice before tamoxifen administration (**Fig. 2A–G, Table S3**). However, following tamoxifen injection, *Prss56^fl/fl^;Glast^Cre^* mice exhibited modest shortening in axial length and VCD, with statistical significance for axial length reached at 4 months (**Fig. 2A-D**). Retinal thickness was also modestly increased at 3 mo and 4 mo, with statistical significance detected at 3 mo but not 4mo (**Fig. 2F**). Consistent with a shortened ocular axial length, hyperopic refraction and a significant increase in retinal *Adamts19* expression were observed in *Prss56^fl/fl^;Glast^Cre^*mice aged to 4 mo (**Fig. 2H–I**).

**Figure 2.**
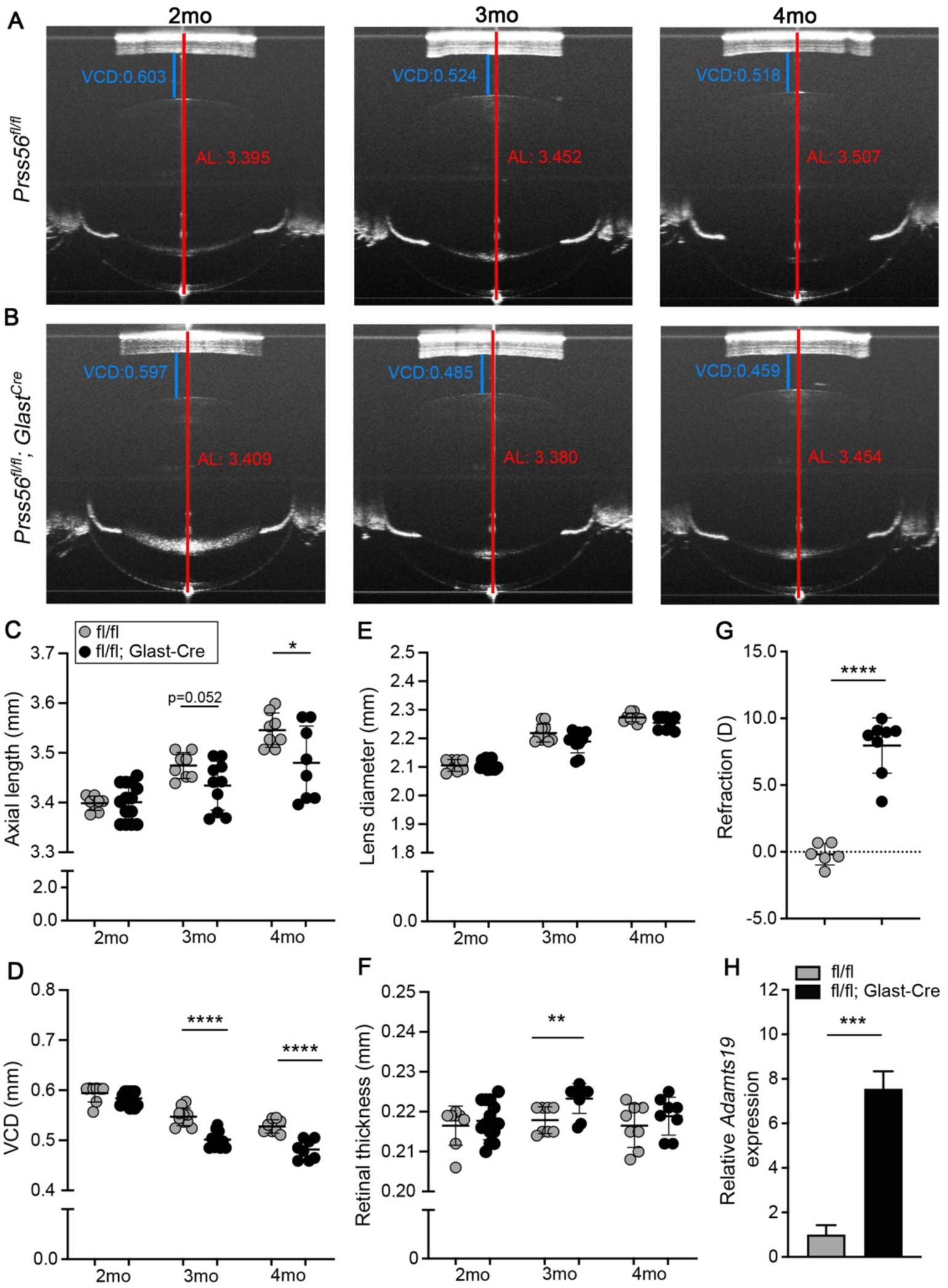
Selective inactivation of *Prss56* in Müller glia at 2 months of age leads to a shortened ocular axial length. *Prss56* was selectively ablated in Müller glia at 2 months (mo) of age by crossing *Prss56^fl/fl^* mice to the inducible *Glast-Cre* mouse strain (*Glast-Cre^ERT^*). Expression of Cre in Müller glial cells was induced by tamoxifen injection at 2mo. (**A-B**) Representative OCT images from 2mo, 3mo and 4mo mice showing reduced axial length and vitreous chamber depth (VCD) at 3mo and 4mo following conditional inactivation of *Prss56* from Müller glia at 2mo (compare eyes from injected *Prss56^fl/fl^; Glast^Cre^* mice in B to those from injected *Prss56^fl/fl^* littermates in A). (**C-F**) Ocular biometric parameters were similar between *Prss56^fl/fl^* and *Prss56^fl/fl^;Glast^Cre^* mice prior to tamoxifen injection at 2mo. Following tamoxifen injection, reductions in ocular axial length (C) and VCD (D) were detected in *Prss56^fl/fl^;Glast^Cre^*mice compared to their *Prss56^fl/fl^* littermates at both ages examined (3mo and 4mo), while an increase in retinal thickness was observed at 3mo and 4mo, statistical significance was only observed at 3mo (F). n=14 *Prss56^fl/fl^; Glast^Cre^* eyes and 8 control *Prss56^fl/fl^* eyes at 2mo; n= 10 *Prss56^fl/fl^; Glast^Cre^* eyes and 8 control *Prss56^fl/fl^* eyes at 3mo; n=8 eyes/group at 4mo. (**G**) Consistent with reduced ocular axial length, *Prss56^fl/fl^; Glast^Cre^* mice injected with tamoxifen at 2mo display hyperopic refraction compared to control *Prss56^fl/fl^* littermates at 4 mo. n= 6 *Prss56^fl/fl^* and 8 *Prss56^fl/fl^; Glast^Cre^* eyes. (**H**) Following tamoxifen injection at 2mo, *Adamts19* expression was also significantly increased in retina from *Prss56^fl/fl^; Glast^Cre^*mice compared to *Prss56^fl/fl^* littermates aged to 4 mo. n= 3 eyes/group. Data are presented as mean ± SD; *p<0.05; ** p<0.01; *** p<0.001; **** p<0.0001, unpaired two-tailed Student t-test. Additional ocular parameter measurements are presented in Supplemental Table 3.

To further define the temporal window of PRSS56 activity in regulating ocular growth, we inactivated *Prss56* in Müller glia at 3 mo, a stage generally considered to be well beyond the emmetropization period.^24^ Interestingly, even at this late time point, *Prss56^fl/fl^;Glast^Cre^*mice showed a measurable shortening of axial length and VCD compared to age-matched controls at 4 mo and 5 mo (**Fig. S3**, **Table S4**). This unexpected result reveals that fine-tuning of ocular axial length continues beyond 3 mo in mice and that PRSS56 remains a key factor in modulating this process. Together, these findings establish that Müller glia-derived *Prss56* is an integral part of the ocular axial length regulation machinery that is required continuously from prenatal life to adulthood in mice.

### PRSS56-driven ocular axial elongation is not dependent on light

To assess whether PRSS56-dependent axial growth is responsive to light-dependent visual cues, *Prss56^fl/fl^;Glast^Cre^* mice and littermate controls were maintained in complete darkness for one month following tamoxifen injection at 1 mo. This dark-rearing paradigm effectively eliminates light-driven visual stimulation, allowing us to investigate the contribution of PRSS56 in the absence of light-dependent cues. Ocular biometric analysis revealed that dark-reared *Prss56^fl/f^;Glast^Cre^*mice exhibit significantly shortened axial length and VCD, along with a modest but consistent increase in retinal thickness relative to control *Prss56^fl/fl^* mice (**Fig. 3, Table S5**) that are reminiscent of ocular changes observed in normally reared, light-exposed cohorts (**Fig. 1**). These results indicate that PRSS56 promotes ocular growth independently of light-evoked visual stimulation and further support a role for PRSS56 as a key retinal factor contributing to the intrinsic ocular growth program.

**Figure 3.**
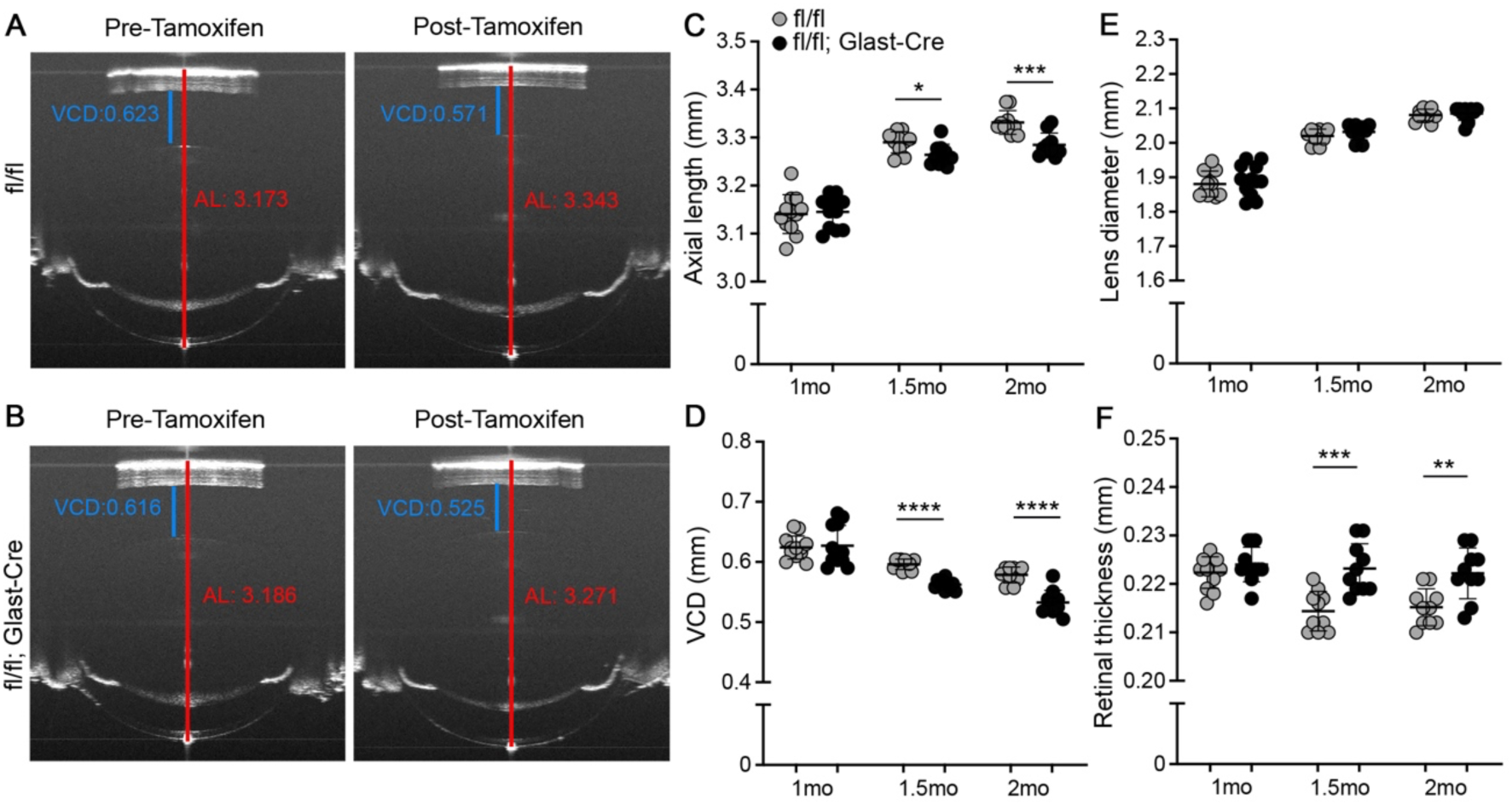
Light is not required for PRSS56-mediated ocular axial elongation following retinal maturation. *Prss56* was selectively ablated from Müller glia at 1 month (mo) of age by crossing *Prss56^fl/fl^* mice to the inducible *Glast-Cre* mouse strain (*Glast-Cre^ERT^*). Following tamoxifen injection at 1mo, mice were reared under dark conditions. (**A-B**) Representative OCT images from 1mo (Pre-Tamoxifen) and 2mo mice (Post-Tamoxifen) mice showing reduced axial length and vitreous chamber depth (VCD) at 2mo in dark reared animals following conditional inactivation of *Prss56* from Müller glia at 1mo (compare eyes from tamoxifen-injected *Prss56^fl/fl^; Glast^Cre^* mice in B to those from injected *Prss56^fl/fl^* littermates in A). (**C-F**) Ocular biometric parameters were similar between *Prss56^fl/fl^* and *Prss56^fl/fl^;Glast^Cre^* mice prior to tamoxifen injection at 1mo. Following tamoxifen administration and dark rearing, modest but significant reductions in ocular axial length (C) and VCD (D) were detected at in *Prss56^fl/fl^;Glast^Cre^*mice compared to their *Prss56^fl/fl^* littermates at 1.5mo and 2mo, while a substantial increase in retinal thickness was observed at 1.5 and 2 mo. n=12 *Prss56^fl/fl^; Glast^Cre^* eyes and 12 control *Prss56^fl/fl^* eyes at 1mo; n=10 *Prss56^fl/fl^; Glast^Cre^* eyes and 10 control *Prss56^fl/fl^* eyes at 1.5mo and 2mo. Data are presented as mean ± SD; *p<0.05; ** p<0.01; *** p<0.001; **** p<0.0001, unpaired two-tailed Student t-test. Additional ocular parameter measurements are presented in Supplemental Table 5.

### AAV-mediated PRSS56 overexpression in Müller glia promotes ocular axial elongation

To further establish the role of PRSS56 in promoting ocular axial growth, we used a reciprocal gain-of-function strategy to selectively overexpress PRSS56 in Müller glia. To this end, we used an adeno-associated virus (AAV2)- mediated gene delivery approach using a Cre-dependent double-floxed inverted *Prss56* open reading frame construct (DIO-*Prss56*). The AAV2-DIO-*Prss56* were intravitreally injected into one eye of *Glast*-*Cre^ERT^*mice (**Fig. S4**), with the uninjected contralateral eye serving as an internal control. In this experimental setup, tamoxifen administration will lead to *Glast*-Cre mediated inversion and overexpression of PRSS56 in Müller glia. Baseline ocular biometry performed before tamoxifen administration at around 1.5 mo confirmed that axial length, VCD, lens diameter, and retinal thickness were comparable between AAV-injected and uninjected control eyes (**Fig. 4A-F, Table S6**). However, about 2 weeks post-tamoxifen induction, the AAV-injected eyes displayed significantly increased axial length and VCD, as well as a reduction in retinal thickness, relative to contralateral uninjected control eyes (**Fig. 4A-D**, **Fig. 4F**). These phenotypic changes were not observed in the AAV2-DIO-*Prss56* injected eyes in the absence of *Glast*-*Cre^ERT^* (**Fig. S5A-F**) and support a gain-of-function effect, whereby elevated PRSS56 levels actively promote ocular elongation that is mainly attributed to an increase in VCD as observed in clinical myopia. Consistent with these changes, *Prss56^fl/fl^;Glast^Cre^*mice also showed myopic refraction (**Fig. 4G**). Intravitreal injection of AAV2 carrying DIO-*mCherry* reporter construct was used to validate *Glast*-*Cre^ERT^*-mediated recombination in Müller glia. Following tamoxifen administration, mCherry signal was specifically detected in Müller glia (**Fig. S5G**). To assess whether PRSS56 overexpression caused any structural or pathological abnormalities, we performed fundoscopy, fluorescein angiography and histological analyses. Despite the observed retinal thinning, retinal vasculature remained intact and appeared normal, and no gross morphological abnormalities observed in the overall ocular globe structure and retina (**Fig. 5**). Thus, retinal PRSS56 overexpression does not overtly compromise ocular morphology or vascular integrity. To determine the time kinetics of PRSS56-induced ocular elongation, we conducted a time course experiment in which axial length and VCD were measured every four days following tamoxifen induction, beginning at day 4 and continuing through day 20 in a *Glast*-*Cre^ERT^* mouse. A significant increase in axial length and VCD were already detectable by day 4 and persisted up to day 20, the final time point examined (**Fig. S6**). These findings demonstrate that PRSS56-driven ocular elongation occurs rapidly and is sustained over time, further supporting Müller glial derived PRSS56 as an autonomous retinal regulator of axial growth in the postnatal eye.

**Figure 4.**
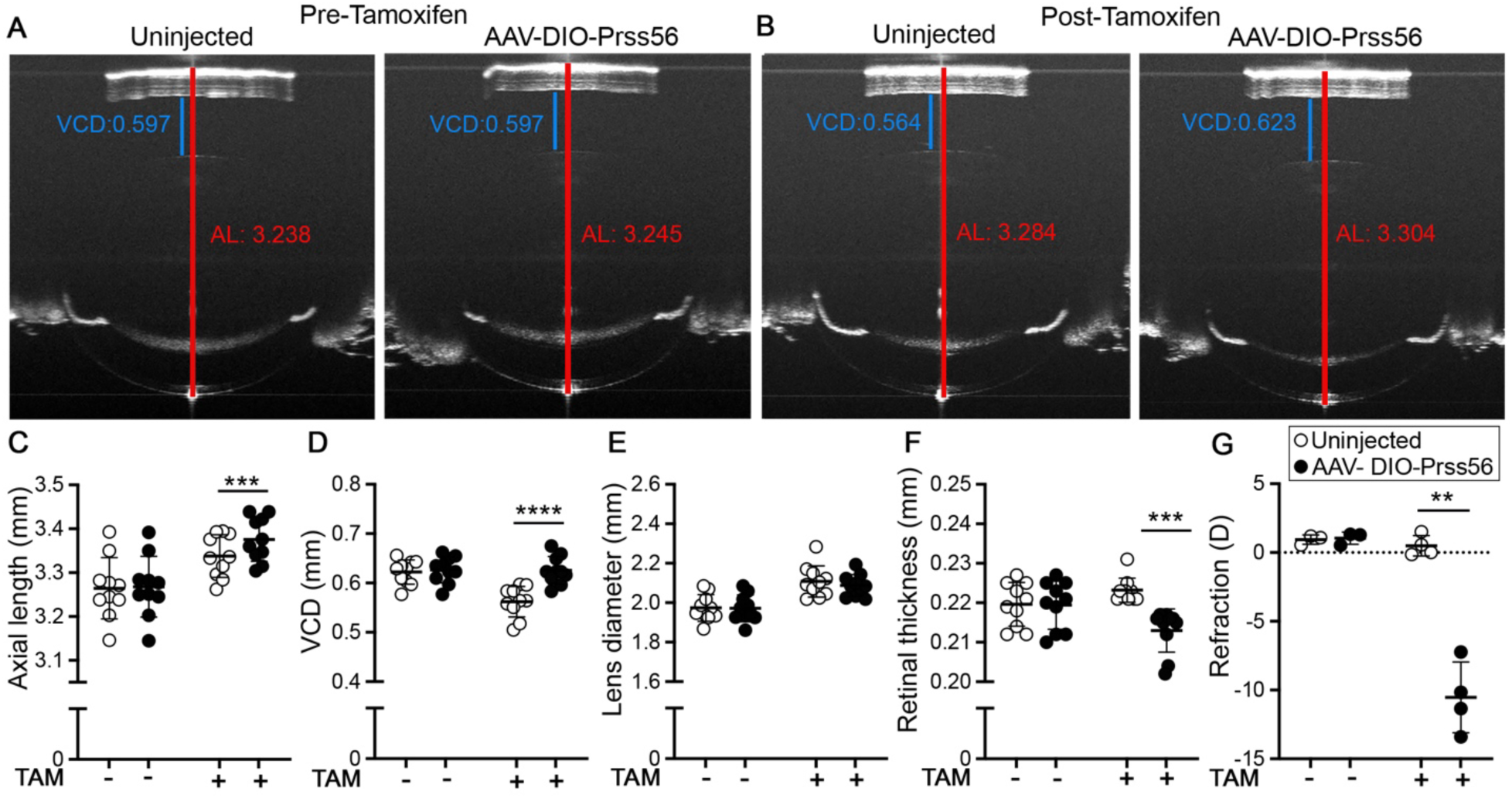
AAV-mediated overexpression of *Prss56* in Müller Glia leads to increased ocular axial elongation. Selective overexpression of *Prss56* in Müller Glia using the AAV-DIO-Prss56 construct in presence of *Glast*-*Cre^ERT^* was induced by tamoxifen administration around 1.5 mo. (**A-B**) Representative OCT images showing that conditional overexpression of PRSS56 in Müller Glia at an age when retina is fully mature (around 1.5 mo of age) lead to significant increases in ocular axial length and vitreous chamber depth (VCD) about 2 weeks after tamoxifen administration (compare AAV-DIO-*Prss56*-injected and contralateral uninjected control eyes in B). (**C-F**) Ocular biometric parameters were similar between AAV-DIO-*Prss56* injected and uninjected control prior to tamoxifen administration around 1.5 mo. Following tamoxifen injection, significant increases in ocular axial length (C) and VCD (D) and reduction in retinal thickness (F) were detected in AAV-DIO-*Prss56* injected eyes compared to contralateral uninjected control eyes at 2-3 mo. n=7 eyes/group. (**G**) Consistent with ocular axial elongation, AAV-DIO-*Prss56* injected eyes exhibited myopic refraction following tamoxifen administration compared to contralateral uninjected control eye at 3 mo. n=3-4 eyes/group.

**Figure 5.**
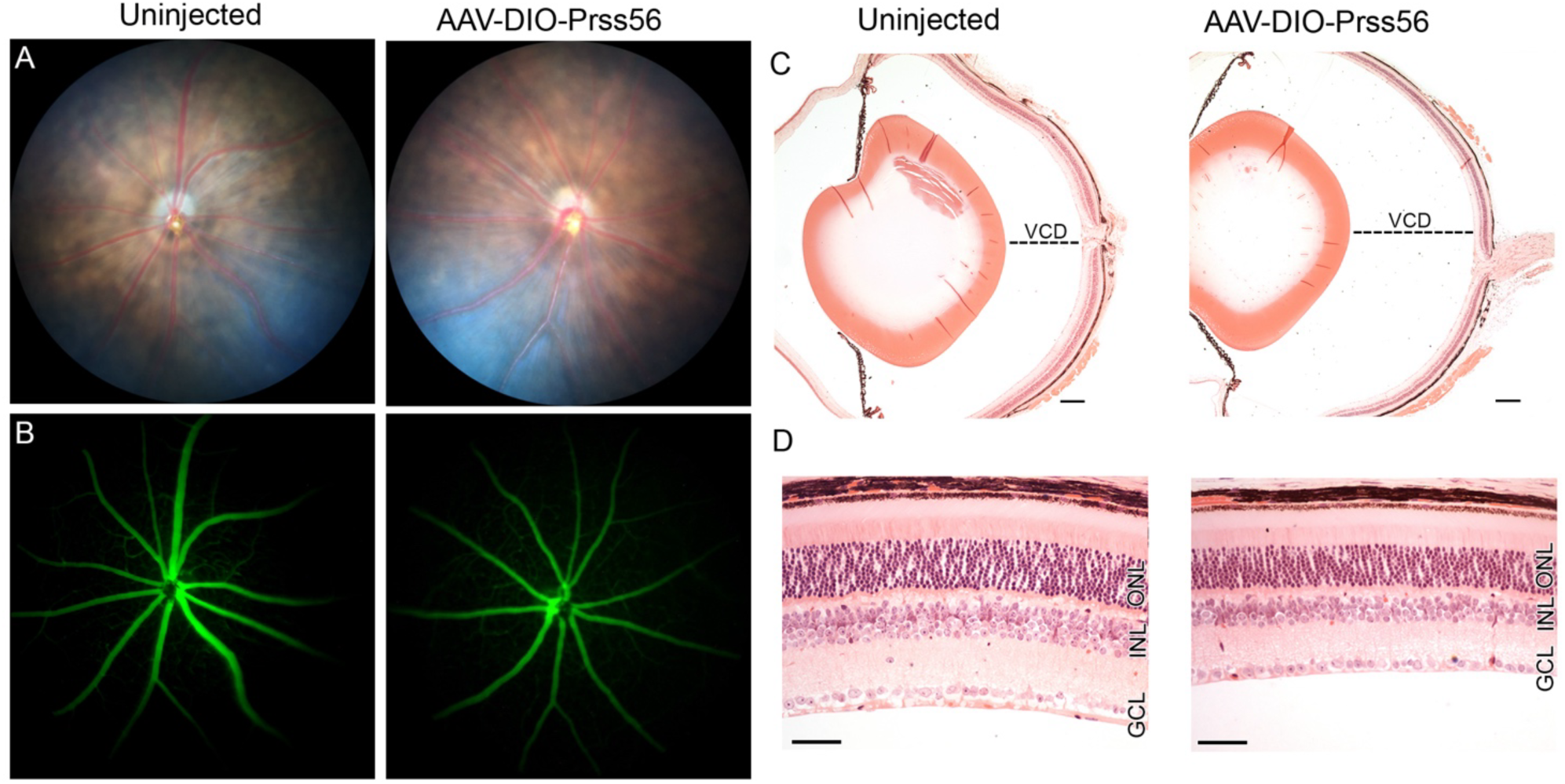
Increased ocular axial elongation resulting from AAV-mediated overexpression of *Prss56* in Müller Glia is accompanied by retinal thinning. Selective overexpression of *Prss56* in Müller Glia using the AAV-DIO-Prss56 construct in presence of *Glast-Cre^ERT^*was induced by tamoxifen administration around 1.5mo. (**A**) Fundoscopy and (**B**) Fluorescein angiography revealed no gross retinal or vascular abnormality in AAV-DIO-Prss56 injected eyes. (**C-D**) Representative H&E-stained ocular sections showing enlarged VCD (C) and retinal thinning (D) in AAV-DIO-Prss56 injected eyes compared to contralateral uninjected control eyes at 3 mo. Scale bars = 100μm (C), 50μm (D). GCL, ganglionic cell layer; INL, inner nuclear layer; ONL, outer nuclear layer.

### PRSS56-mediated ocular axial elongation requires a functional catalytic domain

To determine whether PRSS56-driven ocular axial elongation depends on its proteolytic function, we used an AAV2-DIO construct encoding a catalytically inactive form of PRSS56 (AAV2-DIO-PRSS56-Cat mut). The AAV2-DIO construct was injected intravitreally into *Glast*-*Cre^ERT^* mice as described above. Baseline ocular biometry prior to tamoxifen administration was comparable between AAV-injected and uninjected control eyes. In contrast to AAV-mediated overexpression of wild-type PRSS56, tamoxifen-induced expression of the PRSS56 catalytic mutant failed to induce significant change in axial length or VCD relative to uninjected eyes (**Fig. 6A–D, Table S7**). To confirm proper expression of the mutant protein, HEK293 cells were transduced with an AAV2-DIO vector expressing FLAG-tagged PRSS56-Cat Mut in the presence or absence of Cre recombinase. Western blot analysis with an anti-FLAG antibody confirmed Cre-dependent expression of the catalytic mutant (**Fig. S7A)**. Furthermore, the expression of c-terminus FLAG-tagged PRSS56 catalytic mutant was detected in the retina following intravitreal delivery of AAV2-DIO vector in *Glast-Cre^ERT^* mice and tamoxifen induction (**Fig. S7B)**. Together, these findings demonstrate that PRSS56-mediated ocular axial elongation is dependent on its catalytic activity.

**Figure 6.**
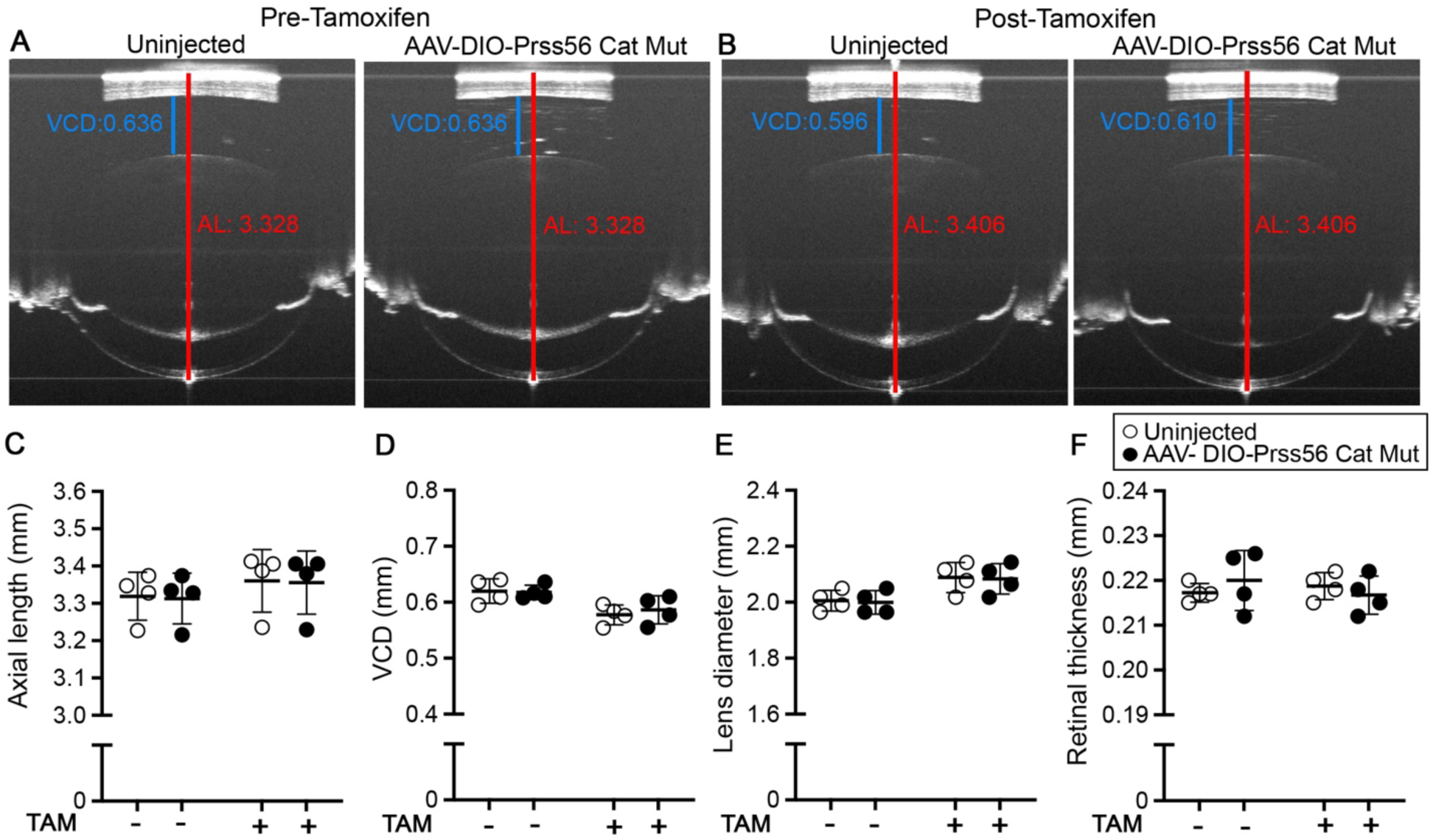
AAV-mediated overexpression of a catalytically inactive form of PRSS56 in Müller Glia does not lead to ocular axial elongation. Selective overexpression of a catalytically inactive form of PRSS56 in Müller Glia using the AAV-DIO-Prss56 Cat-Mut construct in presence of *Glast*-*Cre^ERT^* was induced by tamoxifen administration around 1-1.5 mo. (**A-B**) Representative OCT images showing that conditional overexpression of a catalytically inactive form of PRSS56 in Müller Glia at an age when retina is fully mature (around 1-1.5 mo of age) does not alter ocular axial length and vitreous chamber depth (VCD) about 2 weeks after tamoxifen administration (compare AAV-DIO-Prss56 Cat Mut-injected and contralateral uninjected control eyes in B). (**C-F**) Ocular biometric parameters were similar between AAV-DIO-Prss56 Cat Mut injected and uninjected control prior and after tamoxifen administration. n=4 eyes/group. Additional ocular parameter measurements are presented in Supplemental Table 7.

### PRSS56 variant linked to axial length is associated with myopia correlating with disease severity in the GERA cohort

Common genetic variation at the *PRSS56* locus has previously been associated with axial length (AL) and refractive error.^27,28^ To better define the phenotypic context of this association, we performed GWAS analyses of myopia in the GERA (Genetic Epidemiology Research on Adult Health and Aging) cohort non-Hispanic White (NHW) sample using mean spherical equivalent (MSE) to classify individuals into non-myopic/control (N=36,487), myopic (N=19,540), and high myopic (N=2,806) subgroups.^29^ *PRSS56* variant rs2853447 (which is in strong Linkage Disequilibrium (LD) [r^2^=0.75; D’=87] with lead SNP (Single Nucleotide Polymorphism) rs1550094 previously reported to be associated with MSE)^26^ showed a significant association with all myopia (OR [SE]= 0.91 [0.016]; P=1.59x10^-7^), and an even stronger association with high myopia (OR [SE]= 0.82 [0.035]; P=3.85x10^-9^) **(Fig. S8)**. These analyses confirm *PRSS56* as a myopia-associated risk locus and indicate that its effect size is amplified in individuals with more severe refractive error.

To assess whether the *PRSS56* variant contributes to axial elongation in a refractive error–dependent manner, we next performed genetic association analyses of axial length as a quantitative trait within each refractive subgroup: non-myopic (N = 8,053), all myopic (N = 4,307), and high myopic (N = 689) subgroups (**Fig. S9** and **Table 1 and 2**). Interestingly, while suggestive associations between rs2853447 and axial length were observed in both all-myopic and high-myopic groups (**Fig. S9B, C**), no association was detected in the non-myopic group (**Fig. S9A**). Notably, despite having the smallest sample size, the strongest association was detected in the high-myopic group (**Table 2**).

**Table 1.** Characteristics of GERA participants (all non-Hispanic white individuals) included in the different genetic association analyses

| | **N** | **AL (mm)**<br>Mean ± SD | **MSE (Diopters),**<br>Mean ± SD | **Age at 1<sup>st</sup> AL**<br>**measurement (years),**<br>Mean ± SD |
| --- | --- | --- | --- | --- |
| **Axial length (AL) as**<br>**a quantitative trait** | 16,523 | 24.01 ± 1.26 | -0.30 ± 2.59 | 75.4 ± 8.6 |
| All Myopia | 4,307 | 24.95 ± 1.25 | -3.03 ± 2.20 | 73.38 ± 8.41 |
| High myopia | 689 | 26.33 ± 1.15 | -7.11 ± 1.91 | 70.70 ± 8.23 |
| No myopia | 8,053 | 23.50 ± 0.91 | 1.15 ± 1.27 | 76.47 ± 8.27 |
| **Myopia based on**<br>**MSE (case/control)** | | | | **Age at 1<sup>st</sup> MSE**<br>**measurement (years),**<br>Mean ± SD |
| All Myopia cases | 19,540 | - | -2.83 ± 2.07 | 64.76 ± 12.44 |
| High myopia cases | 2,806 | - | -6.86 ± 1.73 | 60.52 ± 11.44 |
| No myopia controls | 36,487 | - | 0.87 ± 1.19 | 68.83 ± 11.29 |

Abbreviations: N, number of participants; SD, standard deviation; MSE, mean spherical equivalent.

**Table 2.**
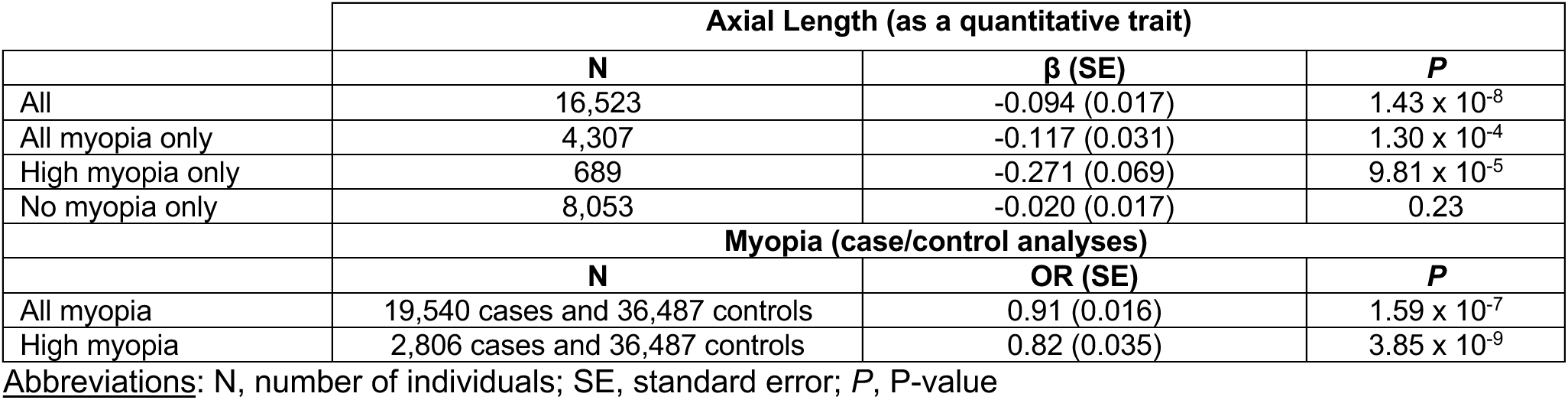
Genetic association results of rs2853447 at *PRSS56* with axial length (as a quantitative trait) and myopia (case/control analyses) in the GERA cohort (non-Hispanic white individuals).

### Myopia- associated PRSS56 risk variants are located within a Cis-Regulatory element

To investigate the potential regulatory function of the *PRSS56* risk variant, we analyzed chromatin accessibility and epigenomic data from human retina.^39^ Notably, intron 4 of the *PRSS56* gene exhibits hallmark features of active cis-regulatory elements. ATAC-seq analysis of whole retina revealed two prominent open chromatin regions at the *PRSS56* locus, one at the proximal promoter and another within intron 4 harboring *PRSS56* variant rs2741297 (which is in strong LD with the lead axial length associated SNP rs2853447). CUT&RUN (**C**leavage **U**nder **T**argets and **R**elease **U**sing **N**uclease) profiling of retinal tissue demonstrates that the intron 4 region with open chromatin overlapped with strong peaks of H3K27ac and H3K4me2, histone modifications associated with active enhancers (**Fig. 7**). These marks indicate that the intron 4 region likely functions as an active regulatory element in the retina. These regions were also enriched for key retinal transcription factors, including OTX2, NRL, and CTCF, a chromatin organizer within the intron 4 region (**Fig. 7**). These transcription factors are known to regulate gene expression in retinal progenitors and photoreceptor lineage specification, suggesting that this intron 4 region may integrate developmental or cell type specific signals to modulate *PRSS56* expression. Additionally, analysis of publicly available single-cell ATAC-seq data from human retina^40^ confirmed the presence of an open chromatin peak at the *PRSS56* promoter and a distinct accessible region at intron 4 (**Fig. S10**). Importantly, these signals based on single-cell RNA-seq from human retina were present in Müller glia, a cell type that endogenously expresses PRSS56 (**Fig. S10**). Together, our genomic findings support a potential role of *PRSS56* variants in modulating PRSS56 expression via cis-regulatory mechanisms, contributing to its role in promoting axial elongation and susceptibility to high myopia.

**Figure 7.**
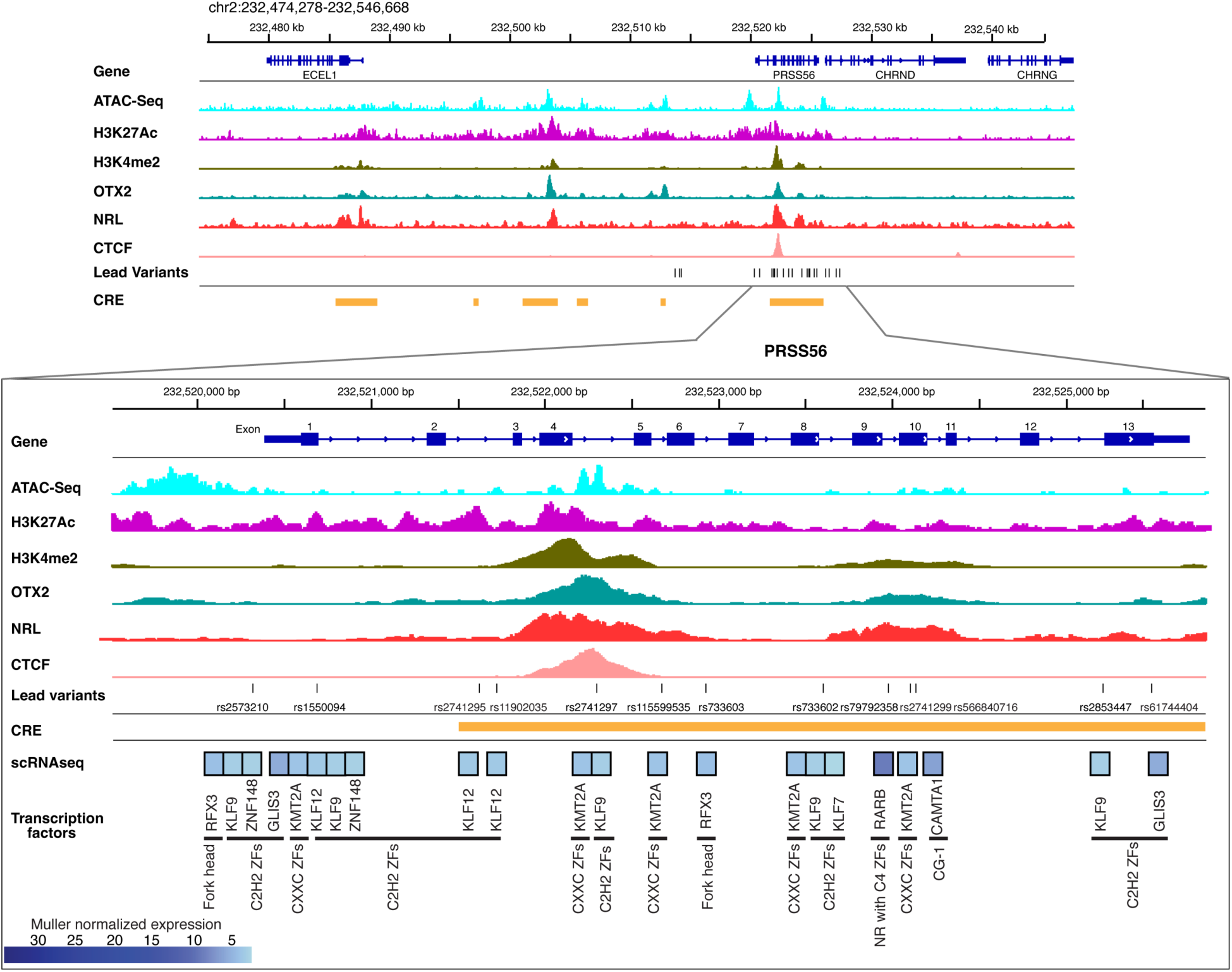
Top panel: Plot showing the enrichment of chromatin accessibility, H3K27ac, H3K4me2, OTX2, NRL, CTCF marks and CREs at the *PRSS56* locus and neighboring genes in the retina. **Bottom panel:** The gene structure of *PRSS56* with enrichment of chromatin accessibility, H3K27ac, H3K4me2, OTX2, NRL, CTCF marks, CRE and lead variants. Selected transcription factor motifs enriched in CREs with overlapping variants and accessible genomic footprints. The accompanying heatmap shows normalized gene expression in Müller cells from single-cell RNA-seq of human retina. Transcription factor families, classified according to TFClass, are indicated below. Abbreviations: CRE, cis-regulatory elements; Fork head, Fork head/winged helix factors; C2H2 ZFs, C2H2 zinc finger factors; CXXC ZFs, CXXC zinc finger factors; NR with C4 ZFs, Nuclear receptors with C4 zinc fingers; CG-1, CG-1 domain factors.

To further explore whether myopia-associated variants in the *PRSS56* locus may affect gene regulation, we examined whether these SNPs are located within predicted transcription factor binding sites. We focused on the genomic interval spanning the entire PRSS56 gene previously associated with refractive error and axial length (**Fig. 7 Bottom Panel**). We further integrated the putative cis-regulatory elements from human retina epigenomics data^39^ and found 11 SNPs mapped to *PRSS56* locus were overlapping the cis-regulatory elements. Within this region, we identified a set of predicted transcription factor binding sites based on sequence motif analysis and regulatory annotations (**Fig. 7 Bottom Panel**, **Table S8**). We then assessed whether the corresponding transcription factors are expressed at appreciable levels in Müller glia. This allowed us to prioritize transcription factors with potential functional relevance in regulating PRSS56 in its endogenous cellular context (**Table S8 and S9**). Several PRSS56-associated SNPs, including rs2741297, were found to overlap or lie near the binding motifs for transcription factors expressed in Müller glia including RORA, KLF9, KLF12, ZNF281, and TCF4 (**Fig. 7 Bottom panel**, **Table S8 and S9**). The proximity of these SNPs to 000000TF motifs suggests they may confer risk by altering transcription factor binding affinity or occupancy, subsequently modulating *PRSS56* expression levels.

## Discussion

In this study, we utilized complementary genetic approaches in the mice in combination with genetic and genomic analyses of human data to establish that PRSS56 is an integral component of the intrinsic growth machinery involved in postnatal ocular axial elongation. We further demonstrate that dysregulation of *PRSS56* transcription contributes to axial length alterations and myopia susceptibility.

Specifically, utilizing conditional gene targeting approach in the mouse we demonstrate that Müller-glia derived PRSS56 is required for ocular elongation at late postnatal developmental stages when retina is matured and emmetropization processes are active in the mouse. Furthermore, these findings reveal that the ocular growth promoting activity of PRSS56 continues beyond the emmetropization period in mice (>3 month), suggesting that axial elongation may continue into adulthood, albeit at a much slower pace. Using a reciprocal gain of function strategy, we show that overexpression of PRSS56 in the mature retina is sufficient to induce an increase in ocular axial length that is accompanied by a concomitant myopic shift in refraction. Further, the failure of the catalytically inactive PRSS56 mutant protein to induce ocular axial elongation demonstrates that its proteolytic activity is essential for PRSS56 function and postnatal eye growth. This supports the notion that PRSS56 drives ocular axial elongation by processing substrate(s) localized in the retina/RPE. These findings suggest that inhibiting PRSS56 catalytic activity may be an effective way to slow excessive axial growth associated with high myopia.

Our results in mice demonstrate that PRSS56 can promote ocular axial growth in the absence of light and is sufficient to drive axial elongation when overexpressed in Müller glia, supporting its role as an intrinsic ocular growth signal. Interestingly, studies in marmoset models have shown that defocus-induced axial elongation is associated with increased retinal *Prss56* mRNA levels, suggesting that visual experience likely modulates *Prss56* expression.^35^ Collectively, these findings support the notion that besides its role in promoting intrinsic (vision-unadjusted) ocular growth, PRSS56 may also serve as a downstream effector of vision-dependent axial growth, translating visual input into molecular signals promoting ocular elongation during emmetropization. Ocular axial elongation is thought to be driven by a cascade of signaling events originating from the retina, transmitted through the retinal pigment epithelium (RPE), choroid, and ultimately resulting in scleral ECM remodeling.^16,17^ Since PRSS56 is predominantly expressed in Müller glia, a retinal cell type that spans the entire thickness of the retina it is ideally suited to coordinate signal transmission across retinal layers and potentially the RPE. Our previous findings suggest that PRSS56 mediated retinal signals are passed on to the RPE to promote ocular axial elongation.^34^ Thus, it is possible that PRSS56 is part of a multi-cellular growth signaling cascade, integrating inputs from retinal neurons and directing downstream changes in the sclera through the RPE.

Complementing the findings from our mouse studies, we performed a stratified analysis of a previously published human genetic study showing an association between *PRSS56* variants and axial length.^29^ No association was observed between *PRSS56* variants and axial length changes in the nonmyopic group, despite its large sample size. In contrast, suggestive associations were detected in the myopic and high myopic groups, with the strongest signal observed in the high myopia subgroup, which had the smallest sample size. The disproportionately strong signal in the high myopia subgroup suggests that the effect of the lead *PRSS56* SNP, rs2853447 on axial length is more penetrant in eyes already predisposed to myopia and that *PRSS56* variants likely act in concert with additional genetic and/or environmental factors to drive excessive ocular elongation and myopia progression (**Fig. 8**).

**Figure 8.**
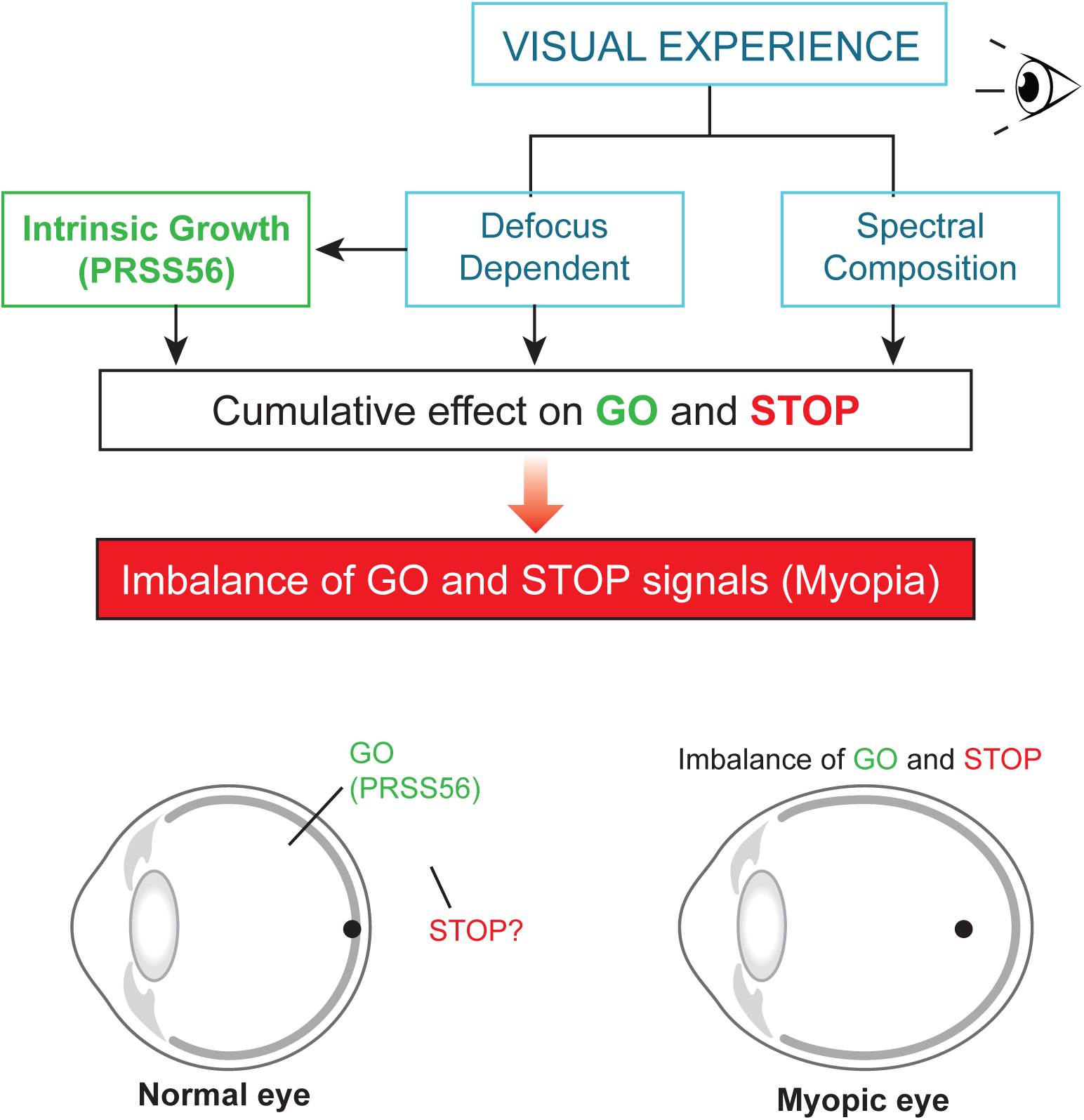
Framework illustrating a unified model integrating intrinsic and visual experience–driven pathways that regulate ocular axial growth and myopia susceptibility. Top panel: Postnatal ocular growth is governed by the concerted action of an intrinsic, genetically programmed *PRSS56*-mediated pathway and vision-dependent mechanisms such as optical defocus and spectral composition dependent signaling. These pathways act together to balance growth-promoting (GO) and growth-limiting (STOP) signals, ensuring optimal axial length and clear vision. Disruption of this balance, due to genetic predisposition or environmental triggers (e.g., reduced exposure to outdoor light), drives excessive axial elongation and increases myopia risk. **Bottom panel:** For instance, in individuals carrying common *PRSS56* variants associated with myopia, elevated *PRSS56* expression may enhance intrinsic growth signaling. When combined with environmental or genetic influences that weaken STOP signals, this imbalance promotes *PRSS56* activity, resulting in excessive axial elongation and high myopia. This framework highlights how genetic and environmental factors converge on shared growth-regulating mechanisms underlying the multifactorial basis of myopia.

Consistent with findings in mice, our functional genomic analysis supports that common non-coding *PRSS56* variants contribute to myopia susceptibility likely by modulating *PRSS56* expression in a context-dependent manner. Interestingly, the lead *PRSS56* variant rs2853447 functions as an expression quantitative trait locus (eQTL), with the risk (minor) allele A associated with increased *PRSS56* expression in human brain tissue based on GTEx data.^41^ Additionally, the *PRSS56* variant rs2741297 (in strong LD with rs2853447) was found to reside within an open chromatin region in intron 4 of *PRSS56*, which also carries histone marks characteristic of active enhancers (**Fig. 7**). Furthermore, the *PRSS56* locus contain predicted transcription factor binding sites relevant to retinal biology (**Table S9**).^40^ Our motif analysis revealed that several transcription factor binding motifs overlap with PRSS56-associated regulatory regions and potentially responsive to transcriptional factors expressed in Müller glia, including RORA, KLF9, KLF12, ZNF281, and TCF4 (**Tables S8 and S9**). Notably, common variants within or near the genes encoding these transcription factors have also been linked to myopia risk.^26^ For instance, a myopia-associated variant (rs12001208) in a long noncoding RNA located adjacent to the gene coding for KLF9, a transcription factor with known roles in neural differentiation and retinal development.^42^ ^43^ This spatial proximity suggests that variation at this locus could influence KLF9 expression or function, thereby indirectly affecting downstream targets such as PRSS56. Notably, the PRSS56 variant rs2741297 resides near a predicted binding site for KLF9 (**Fig. 7**). These observations raise the possibility that PRSS56 expression may be modulated by a network of transcription factors active in Müller glia, and that genetic variation in this network could contribute to altered expression of PRSS56 and excessive ocular elongation in susceptible individuals. Future studies will experimentally validate the putative cis-regulatory elements that govern PRSS56 expression in Müller glia and assess whether the disease-associated variant disrupts transcription factor binding or regulatory activity. Thus, providing a mechanistic link between noncoding variants, altered PRSS56 expression, and axial elongation.

During vision-guided ocular growth, the eye relies on an active feedback system in which "GO" and "STOP" signals regulates the rate of ocular axial elongation in response to visual defocus.^15,44^ Our genetic and experimental data support a role for PRSS56 as a retinal GO signal that promotes axial elongation and is required both during early developmental stages and later periods when ocular axial growth becomes visually regulated. We propose a model in which intrinsic growth signals such as PRSS56 act in parallel with vision-dependent pathways to regulate ocular size (**Fig. 8**). Under normal conditions, this growth-promoting activity is likely balanced by the growth-limiting STOP signals that help maintain emmetropia. In the context of myopia, especially high myopia, this equilibrium may be disrupted through gene-environment interactions. Variants that increase PRSS56 expression can shift the system toward continued elongation, while environmental factors associated with myopia, such as reduced outdoor exposure, may suppress STOP signaling. Although the precise molecular identity of the STOP signal remains unknown, our results suggest that the interaction between PRSS56 and these counterbalancing pathways may be critical in mediating the impact of *PRSS56* variants and their catalytic activity shared on myopia susceptibility. This may explain why *PRSS56* variants enhance axial growth primarily in the context of myopia, where reduced STOP signaling creates a permissive environment for axial elongation, but have little effect in emmetropic eyes **(Fig. 8)**.

In summary, our findings identify PRSS56 as a key component of an intrinsic retinal pathway that sustains ocular axial growth from early development into adulthood in mice. Overexpression of PRSS56 in Müller glia alone is sufficient to drive axial elongation in a catalytic activity-dependent manner, demonstrating its role as an autonomous retinal growth signal. Human genetic analyses further link PRSS56 variants associated with axial length to increased risk of high myopia, likely through regulatory changes that modulate its expression in Müller glia. The study provides a framework on how genetic and environmental factors may converge on a shared pathway to control eye growth highlighting the complex multifactorial nature of myopia development. To better understand PRSS56’s role in ocular axial elongation, future work will focus on identifying factors involved in the regulatory “STOP” pathway(s) that counterbalance its GO activity and determine the molecular substrates through which its proteolytic activity drives elongation.

## Methods

### Animals

All animal procedures adhered to the Association for Research in Vision and Ophthalmology’s (ARVO) statement on the use of animals in ophthalmic research. Mouse studies were performed in compliance with protocols approved by the Institutional Animal Care and Use Committee (IACUC) at the University of California, San Francisco (#AN153083 and #AN196253). Mice were given access to food and water ad libitum and housed under controlled conditions, including a 12-h light/dark cycle (unless otherwise specified for dark-rearing experiments described below) with ad libitum access to food and water, in accordance with the National Institutes of Health guidelines (NIH). All animal procedures were designed to minimize the number of animals used and their suffering.

### Mouse lines

Three mouse lines were used in this study, and all the lines were backcrossed and maintained on a C57BL/6 background for at least 7 generations. The *Prss56* conditional mutant line (*Prss56^fl^*) carries loxP sites flanking exons 2 to 4 of the *Prss56* gene, allowing Cre-mediated excision and inactivation of PRSS56 protease activity. We have previously validated *Prss56^fl^* mice, which allows conditional gene inactivation through Cre recombinase–mediated excision.^33^ For inducible gene inactivation, *Prss56^fl/fl^* mice were crossed to tamoxifen-inducible Cre driver lines. In the progeny, the Cre activity was induced temporally at defined postnatal stages by intraperitoneal administration of tamoxifen.^33^ Previous studies have shown that heterozygous *Prss56^fl/+^* mice carrying universal Cre recombinase exhibit an axial length phenotype indistinguishable from wild type.^33^ Therefore, heterozygous mice were not included in this study. Here, we used Glast-Cre^ERT^ line [STOCK Tg(Slc1a3-cre/ERT)1Nat/J, Jax Strain #:012586] that enables tamoxifen-inducible Cre expression under the Glast promoter.^45^ In addition, we used the Ubc-Cre^ERT2^ line (C57BL/6.Cg-Tg(UBC-Cre/ERT2)1Ejb) that drives tamoxifen-inducible Cre expression under the human ubiquitin C promoter.^46^ To assess Cre-dependent recombination efficiency and potential leakiness, Cre driver lines were crossed to Rosa26-tdTomato mice and recombination was evaluated in tamoxifen-treated and vehicle-treated littermates. Furthermore, control cohorts were included to evaluate recombination in the absence of tamoxifen and to verify allele-specific excision post induction. Male and female littermates were used for all experiments. Genotyping was performed on genomic DNA extracted from tail biopsies using Proteinase K digestion, followed by PCR with primers specific for the *Prss56* floxed allele and *Cre* (**Table S10**).^33^

### Tamoxifen injection

Tamoxifen (T-5648, Sigma) was dissolved in ethanol (200 mg/ml) and diluted in corn oil to a final concentration of 20 mg/ml. Mice received two intraperitoneal injections of tamoxifen (50 µl per injection) on consecutive days. To inactivate *Prss56* at 1, 2 and 3 months, Cre activation was induced by first tamoxifen injection at P29, P60 and P90, respectively.

### Ocular biometry

Axial length and other biometric parameters, including axial length, vitreous chamber depth (VCD), anterior chamber depth (ACD), lens thickness, and retinal thickness were measured by spectral domain optical coherence tomography (SD-OCT; Envisu R4300, Leica/Bioptigen 4300) as described previously.^33,34^ Mice were anesthetized with ketamine and xylazine, 100 mg/kg, and 5mg/kg, respectively) before imaging. The eyes were first applied with 1% tropicamide ophthalmic solution (Somerset) to dilate the pupil for few minutes. Later the tropicamide was wiped, rinsed with a drop of GenTeal tears (Alcon, Fort Worth, TX) post that the eyes were hydrated with GenTeal gel (Alcon, Fort Worth, TX). The mice were then placed on a platform and aligned to center the Purkinje lines for consistent measurements. Axial length was defined from the corneal surface to the retinal pigment epithelium/choroid boundary. Measurements were taken from both eyes OD (Oculus Dexter) and OS (Oculus Sinister). Following imaging, Antisedan (atipamezole hydrochloride from Zoetis) was injected to reverse the sedative and analgesic effect of anesthesia. Since body weight can influence ocular size, efforts were made to minimize the difference in body weight between experimental and corresponding control groups.

### Refraction

Cycloplegic refraction was measured using an automated infrared photorefractor as previously.^47^ Mice were treated with cyclopentolate to temporarily inhibit lens accommodation and placed facing the photorefractor. Only measurements with centered Purkinje images were accepted. At least 30 to 50 valid readings were averaged per eye. Refraction was assessed bilaterally in all animals.

### Dark rearing

For dark rearing, the *Prss56^fl/fl^;Glast^Cre^* and control *Prss56^fl/fl^* (no cre) mice were given two successive intra-peritoneal tamoxifen injections at P29 & P30, one set of the mice was placed in complete darkness for 30 days and the other set of mice maintained on a 12-h light/dark cycle. Both pre- and post-tamoxifen SD-OCT values were captured and after 30 days dark -reared mice were placed back on regular 12-h light/dark cycle.

### Adeno associated virus (AAV2)-mediated *Prss56* overexpression

AAV2 vectors harboring a <u>D</u>ouble-floxed Inverted <u>O</u>rientation (DIO) construct containing the coding sequence for mouse *Prss56* and its catalytically inactive mutant were constructed and packaged by VectorBuilder (pAAV[FLEXon]-CAG>LL:rev(mPrss56 [NM_027084.2]:rev(LL)). To generate a catalytically inactive form of PRSS56 (Cat Mut), the catalytic triad residues His149, Asp195, and Ser290 were replaced with alanine. The construct features a CAG promoter, paired LoxP and Lox2272 sites, and *Prss56* ORF in reverse orientation (**Fig. S4**). Viral titer was quantified by qPCR targeting AAV2 inverted terminal repeats (ITR) sequences. *Glast-Cre^ERT^* enables inducible, Müller glia–biased recombination in the adult retina, as GLAST is robustly expressed in Müller glia postnatally. Tamoxifen induction in adult mice minimizes developmental recombination and restricts astrocyte labeling. For intravitreal injection, about 1 microliter of AAV2-DIO-Prss56 or AAV2-DIO-Prss56 Cat-Mut (10^13^ vital titer) was administered to one eye of C57BL/6J mice carrying the *Glast-Cre^ERT^*transgene or control C57BL/6J mice at postnatal day 21 (P21), while the contralateral eye was not injected to serve as an internal control. At P42, tamoxifen was delivered systemically (intraperitoneally) to drive Cre recombinase activity in retinal Müller glia, enabling inversion of the *Prss56* cassette and selective PRSS56 expression in retinal Müller Glia. SD-OCT values were captured both pre-tamoxifen (P40) and post-tamoxifen (around 2-weeks post tamoxifen treatment). AAV2 carrying DIO-mCherry (10^12^ vital titer, Viral Vector Core, University of Iowa) was used to confirm largely Müller glial specific gene expression in a *Glast*-Cre dependent manner using the same procedure as described above. Additionally, we maintained no Cre controls where in the absence of Cre, AAV-DIO-Prss56 injection did not alter ocular biometric parameters (**Fig. S5**).

### Fundus Imaging and Fluorescein Angiography

Fundus examination and fluorescein angiography were performed using the Micron IV retinal imaging system (Phoenix Research Labs, Pleasanton, CA), which enables high-resolution imaging of the mouse retina. Mice were anesthetized with isoflurane (induction at 3–4%, maintenance at 1.5–2% in oxygen), and pupils were dilated using 1% tropicamide. The animal was positioned on a heated platform, and Genteal gel (Alcon, Fort Worth, TX) was applied to the cornea to maintain clarity and avoid drying during imaging. For fundus imaging, the eye was carefully aligned with the camera, and brightfield images were acquired using standardized illumination and exposure settings. The optic nerve head, retinal vasculature, and overall retinal morphology were documented.

For fluorescein angiography, 10% sodium fluorescein was injected intraperitoneally (10 µl/g body weight). Retinal images were then collected at sequential time points (typically 1, 3, 5, and 10 minutes post-injection) to evaluate vascular filling and leakage. Fluorescence images were analyzed for abnormalities in vessel appearance or signs of dye extravasation. All imaging was conducted under consistent conditions across experimental groups.

### Quantitative RT-PCR

Retinas were isolated and total RNA extracted using the RNeasy Mini Kit (Qiagen), including on-column DNase treatment. cDNA was synthesized from 1 µg of total RNA using the iScript cDNA synthesis kit (Bio-Rad). Quantitative PCR (qPCR) was performed on a Bio-Rad CFX96 Real-Time Detection System using SYBR Green Supermix and gene-specific primers for *Adamts19*, as previously.^34^ Reactions contained 25 ng cDNA and 0.25 µM primers in a final volume of 10 µl. Cycling conditions were 95°C for 5 s, followed by 60°C for 25 s. Technical duplicates and at least three biological replicates per group were included. Gene expression was normalized to *Actb*.

### Immunofluorescence

Eyes were enucleated and immersion-fixed in 4% paraformaldehyde (PFA) at 4°C overnight, cryoprotected in 30% sucrose O/N, embedded in optimal cutting temperature (OCT) compound, and cryosectioned (10 µm). Sections were mounted using Mowiol containing DAPI for nuclear labeling.

### Histology

For histological analysis, eyes were enucleated and fixed in a solution of 1% paraformaldehyde and 2% glutaraldehyde in 0.1 M sodium cacodylate buffer (pH 7.4). Tissues were then dehydrated and embedded in glycol methacrylate resin. Serial sagittal sections (2 µm thick) through the optic nerve head were cut and stained with hematoxylin and eosin for morphological assessment.^33,48^

### Microscopy

Fluorescence images were acquired using either an AxioImager M1 epifluorescence microscope or an LSM700 laser scanning confocal microscope (Zeiss). Image capture and processing were performed using the manufacturers’ respective software packages.

### Western blot analysis

HEK293T cells were transduced with an AAV2-DIO construct expressing FLAG-tagged PRSS56 (Cat Mut) in the presence or absence of AAV-Cre. Cells were lysed in RIPA buffer, and protein extracts were resolved by SDS–PAGE. FLAG-tagged PRSS56 was detected by Western blotting using a monoclonal anti-FLAG M2 antibody (Sigma).

### Human genetic analyses

The Genetic Epidemiology Research on Adult Health and Aging (GERA) cohort comprises 110,266 adult men and women who are members of the Kaiser Permanente Medical Care Plan, Northern California Region (KPNC). KPNC biometry measurements, including axial length (AL), are collected using the Haag-Streit Lenstar 900 device. GERA participants included in the GWAS of AL analyses had at least one recorded AL measurement on both eyes during the same visit. The mean (in millimeters) AL of an individual’s two eyes was used for the analyses.

As previously described ^26,49,29^, GERA participants included in the GWAS analyses of myopia had at least one spherical equivalent value measured during routine eye examinations. Spherical equivalent was calculated as the sphere + (cylinder/2). The spherical equivalent was selected from the first documented refraction assessment, and the mean of both eyes was used.

GWAS analyses of myopia were performed in the GERA cohort non-Hispanic White (NHW) sample using mean spherical equivalent (MSE) to classify individuals into myopic (any myopia), high myopic, and emmetropic (non-myopic control) subgroups (with myopia cases defined as having a MSE ≤ -0.75 diopters (D), high myopia defined as MSE < -5.0 D, and controls as having a MSE > -0.75 D). GWAS analyses were performed using PLINK v1.90^50^ and adjusting for the following covariates: age, sex, and ancestry principal components (PCs), as previously described .^26,29^ All study procedures were approved by the Institutional Review Board of the Kaiser Foundation Research Institute.

### Human genomic analyses

*Identification of potential transcription factors in regulating PRSS56 expression*: To identify transcription factors (TFs) that may regulate PRSS56, we performed a comprehensive in silico motif enrichment analysis using the full genomic sequence of the human *PRSS56* gene, including 1000 base pairs upstream of the annotated transcription start site to encompass the proximal promoter region. This extended sequence was used as input for a motif discovery tool capable of scanning for known transcription factor binding sites across the input sequence. Motif matches were evaluated against a reference transcription factor database, and statistical enrichment was assessed using a false discovery rate (FDR) threshold. The results were filtered based on FDR < 0.05 (see **Table S8**: q_value_0.05).

*Prioritization of candidate TFs in Müller glia*: To prioritize transcription factors relevant to retinal biology, we integrated motif enrichment data with gene expression data from both bulk and single-cell RNA sequencing datasets of the human retina. Expression levels of candidate TFs (from the FDR < 0.05 list) were cross-referenced against publicly available bulk RNA-seq datasets derived from human control retina samples (see **Table S8**: 0.05_genes_bulk_RNAseq), as well as against a high-resolution single-cell RNA-seq atlas to determine their expression across different retinal cell types. These integrative analyses enabled the prioritization of candidate TFs for downstream validation based on both motif enrichment and retinal cell-type specific expression profiles.

*Overlap of SNPs with regulatory elements:* The myopia associated SNPs in the PRSS56 gene were overlapped with human retina cis-regulatory elements to detect the presence and absence of SNPs within the cis-regulatory element using the closestBed command from bedtools.^51^

*Transcription factor binding analysis*: Based on human retina ATAC-seq data for PRSS56 gene we identified binding motifs present within these footprints at an FDR < 0.05 using FIMO^52^ with the motif database HOCOMOCOv13.^53^ Motifs from expressed transcription factors were filtered to those expressed in a previous retina transcriptome study with >=1 CPM (**Table S8)**.^54^ The expression of these transcription factors was further checked in single cell RNAseq data from human retina and transcription factors with average log expression > 1 in Müller cells was further considered (**Table S8)**.^40^

### Statistical analysis

All Statistical analyses were performed using GraphPad Prism software. Data are presented as mean ± SEM unless otherwise noted. Two group comparisons were performed using an unpaired two-tailed Student test, except for AAV experiments where a paired two tail t-test was used for pairwise comparison between AAV injected and uninjected contralateral control eye. Comparison across multiple groups were performed using one-way ANOVA. P values <0.05 were considered statistically significant. Exact sample sizes (n) for each experiment are provided in the corresponding figure legends and supplemental tables.

### Material availability statement

All unique/stable reagents and including the *Prss56* conditional mutant mouse line and AAV plasmids generated in this study, are available from the Lead Contact, K. Saidas Nair, upon request. A Material Transfer Agreement is needed to comply with the guidelines and policies of the National Institutes of Health.

## Acknowledgements

Genotyping of the GERA cohort was funded by a grant from the National Institute on Aging, National Institute of Mental Health, and National Institute of Health Common Fund (RC2AG036607). HC was supported in part by the National Eye Institute (NEI) RO1EY027004, and RO1EY033010. KSN was supported by the NEI R01EY032666, R01EY033015, Core Grant P30 EY002162, Marin Community Foundation-Masneri Fund, The RPB unrestricted grant, and All May See Foundation. A part of this work was supported by Intramural Research Program of the National Eye Institute (ZIAEY000450 and ZIAEY000546) and utilized the computational resources of the NIH HPC Biowulf cluster (https://hpc.nih.gov). The contributions of the NIH authors are considered works of the United States Government. The findings and conclusions presented in this paper are those of the authors and do not necessarily reflect the views of the NIH or the U.S. Department of Health and Human Services.

## Author Contributions

KSN conceived and supervised the study. K.G. performed the mouse experiments, including ocular biometric measurements and data analysis. D.D. and E.B.U. assisted with animal husbandry, and performed genotyping, qPCR, and Western blot analyses. C.L.D. led the quantitative and statistical analyses. S.K. contributed to the initial mouse studies. Y-M.K. and K.S.N. performed the histological analyses. Y.I. assisted with cell culture experiments and provided guidance on protein studies. J.A., N.S., and A.S. conducted the genomic and transcriptional regulatory analyses. C.J. and H.C. contributed to the human genetic studies. C.L.D. and K.S.N. wrote the manuscript. K.G., D.D., E.B.U., S.K., H.C, J.A., N.S. and A.S. contributed and reviewed the paper. A.S. and H.C. oversaw the genomics and human genetics components of the study.

