## Supplemental Figures for "PRSS56 acts as an intrinsic retinal signal driving postnatal ocular axial growth and myopia susceptibility"

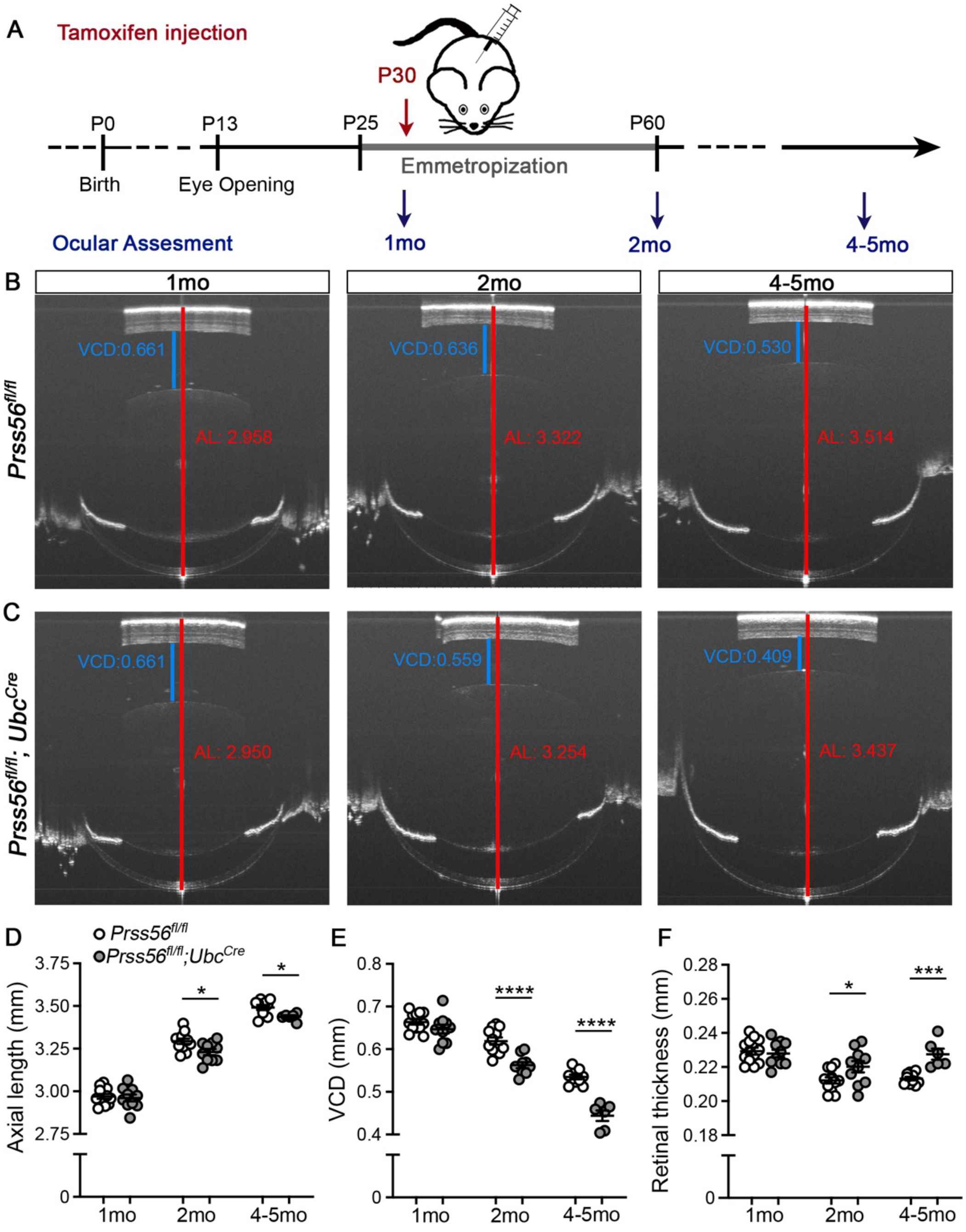


**Supplemental Figure 1. Conditional *Ubc*-Cre-mediated inactivation of *Prss56* at 1 month of age leads to shortened ocular axial length.**

*Prss56* was conditionally ablated in a time-specific manner by crossing *Prss56*$^{fl/fl}$ mice to the inducible ubiquitous *Ubc-Cre* mouse strain (*Ubc-Cre*$^{ERT2}$). (A) Diagram illustrating the timeline of tamoxifen injection and ocular assessment. Cre expression was induced by tamoxifen injection at 1 month (mo) and ocular parameters were measured prior to tamoxifen injection at 1 mo and at two time points post-injection (2 mo and 4-5 mo). (B-C) Representative OCT images prior (P28) and after (2mo and 4-5mo) tamoxifen injection showing reduced axial length and vitreous chamber depth (VCD) following conditional ablation of *Prss56* (compare *Prss56*$^{fl/fl}$ eyes in B to *Prss56*$^{fl/fl}$*;Ubc*$^{Cre}$ eyes in C). (D-F) Ocular biometric parameters were similar between *Prss56*$^{fl/fl}$

and *Prss56*$^{fl/fl}$*;Ubc*$^{Cre}$ mice at 1mo. Following tamoxifen injection at 1mo, significant reductions in ocular axial length (D) and VCD (E) and increase in retinal thickness (F) were detected at in *Prss56*$^{fl/fl}$*;Ubc*$^{Cre}$ mice compared to their *Prss56*$^{fl/fl}$ littermates at 2mo and 4-5mo. n= 12 *Prss56*$^{fl/fl}$ and 10 *Prss56*$^{fl/fl}$*;Ubc*$^{Cre}$ eyes at 1mo and 2mo, and n= 10 *Prss56*$^{fl/fl}$ and 6 *Prss56*$^{fl/fl}$*;Ubc*$^{Cre}$ eyes at 4-5 months. Data are presented as mean ± SD; *p<0.05; *** p<0.001; **** p<0.0001, unpaired two-tailed Student t-test. Additional ocular parameter measurements are presented in Supplemental Table 1.

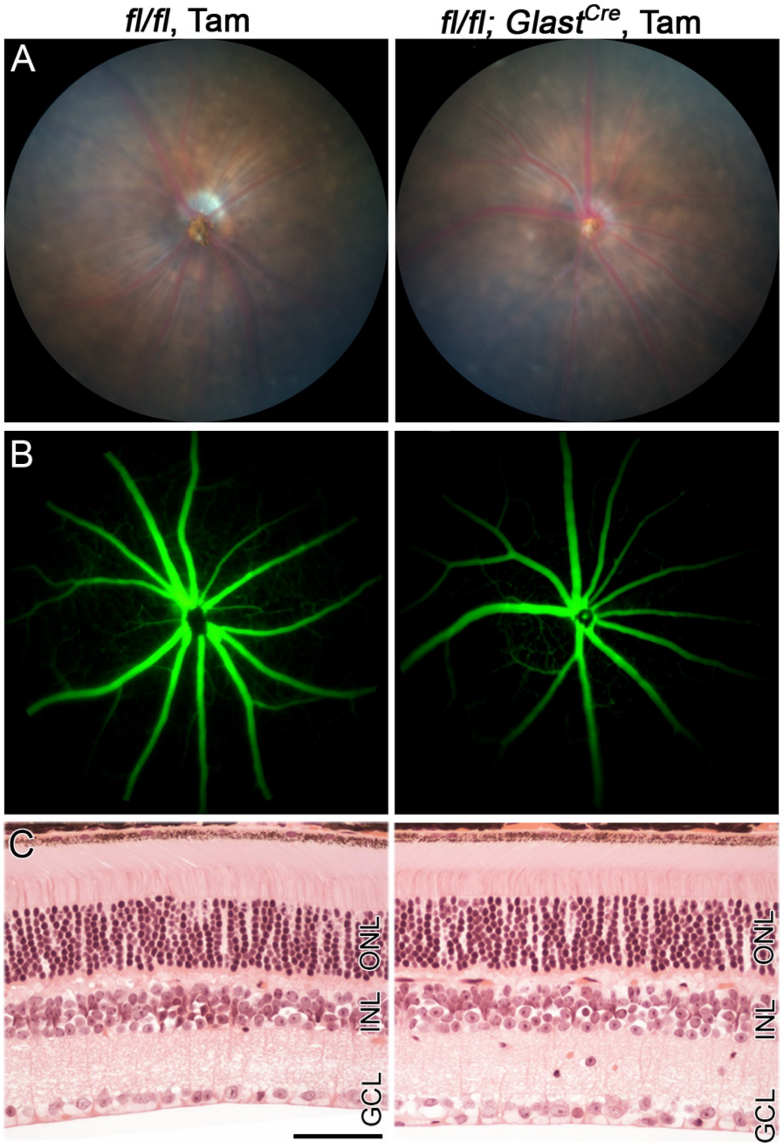


**Supplemental Figure 2. No retinal vascular and morphological abnormalities were observed following selective inactivation of *Prss56* in Müller glia at 1 month of age.**
*Prss56* was selectively ablated in Müller glia at 1 month (mo) of age by crossing *Prss56*$^{fl/fl}$ mice to the inducible *Glast-Cre* mouse strain (*Glast-Cre*$^{ERT}$). Conditional Cre expression in Müller glial cells was induced by tamoxifen injection. (**A**) Funduscopy, (**B**) Fluorescein angiography and (**C**) Histological H&E-staining of ocular sections revealed no gross retinal or vascular abnormality in 3mo *Prss56*$^{fl/fl}$;*Glast*$^{Cre}$ mice following tamoxifen administration. Scale bar = 50μm. GCL, ganglionic cell layer; INL, inner nuclear layer; ONL, outer nuclear layer.

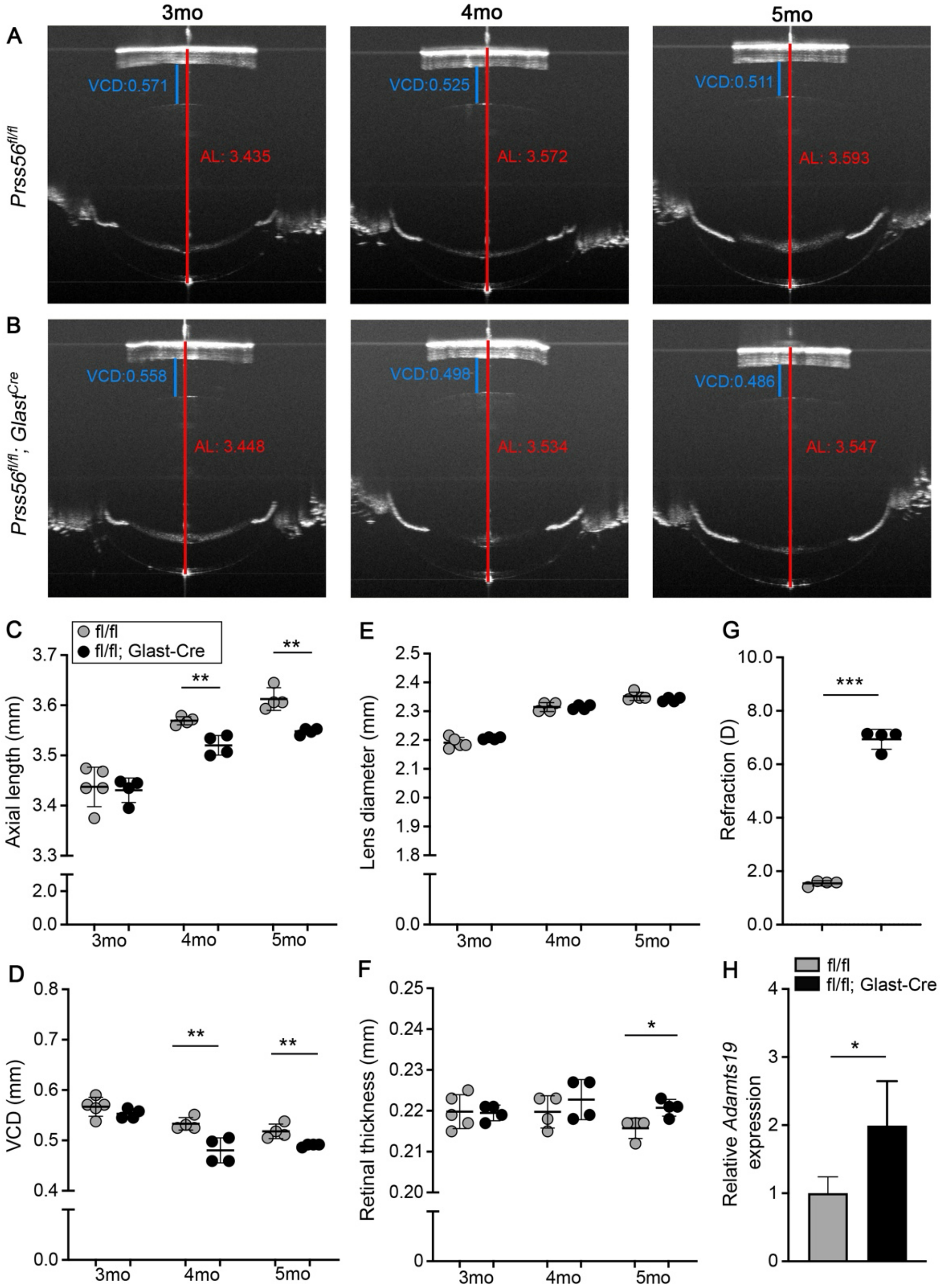


**Supplemental Figure 3. Selective inactivation of *Prss56* in Müller glia at 3 months of age leads to a shortened ocular axial length.**

*Prss56* was selectively ablated in Müller glia at 3 months (mo) of age by crossing *Prss56*$^{fl/fl}$ mice to the inducible *Glast-Cre* mouse strain (*Glast-Cre*$^{ERT}$). Conditional Cre expression in Müller glial cells was induced by tamoxifen injection at 3mo. (**A-B**) Representative OCT images from 3mo,

4mo, and 5mo mice showing reduced axial length and vitreous chamber depth (VCD) at 4mo and 5mo following conditional inactivation of *Prss56* from Müller glia at 3 mo (compare eyes from injected *Prss56*$^{fl/fl}$*; Glast*$^{Cre}$ mice in B to those from injected control *Prss56*$^{fl/fl}$ littermates in A). (**C-F**) Ocular biometric parameters were similar between *Prss56*$^{fl/fl}$ and *Prss56*$^{fl/fl}$*;Glast*$^{Cre}$ mice prior to tamoxifen injection at 3mo. Following tamoxifen injection, reductions in ocular axial length (C) and VCD (D) were detected at in *Prss56*$^{fl/fl}$*;Glast*$^{Cre}$ mice compared to their *Prss56*$^{fl/fl}$ littermates at both ages examined (3mo and 4mo), while a significant increase in retinal thickness was observed by 5mo (F). n=5 *Prss56*$^{fl/fl}$ eyes at 3mo and 4 eyes for each of the other groups. (G) Consistent with reduced ocular axial length, *Prss56*$^{fl/fl}$*; Glast*$^{Cre}$ mice injected with tamoxifen at 3mo display a hyperopic refraction compared to tamoxifen injected control *Prss56*$^{fl/fl}$ littermates at 5 mo. n= 4 *Prss56*$^{fl/fl}$ and 4 *Prss56*$^{fl/fl}$*; Glast*$^{Cre}$ eyes. (**H**) Following tamoxifen injection at 3mo, *Adamts19* expression was also significantly increased in retina from *Prss56*$^{fl/fl}$*; Glast*$^{Cre}$ mice compared to *Prss56*$^{fl/fl}$ littermates at 5 mo. n= 4 *Prss56*$^{fl/fl}$ and 4 *Prss56*$^{fl/fl}$*;Glast*$^{Cre}$ eyes. Data are presented as mean ± SD; *p<0.05; ** p<0.01; *** p<0.001, unpaired two-tailed Student t-test. Additional ocular parameter measurements are presented in Supplemental Table 5.

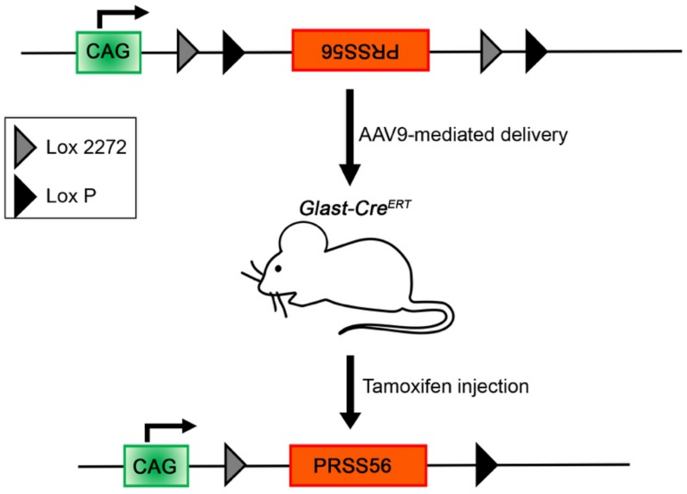


**Supplemental Figure 4. Diagram illustrating the strategy for selective overexpression of PRSS56 in Müller Glia.** Selective overexpression of *Prss56* in Müller Glia was achieved using ocular injection of the AAV-DIO-Prss56 construct in *Glast-Cre*$^{ERT}$ mice to allow for tamoxifen-inducible PRSS56 expression in Müller glial cells at specific time points.

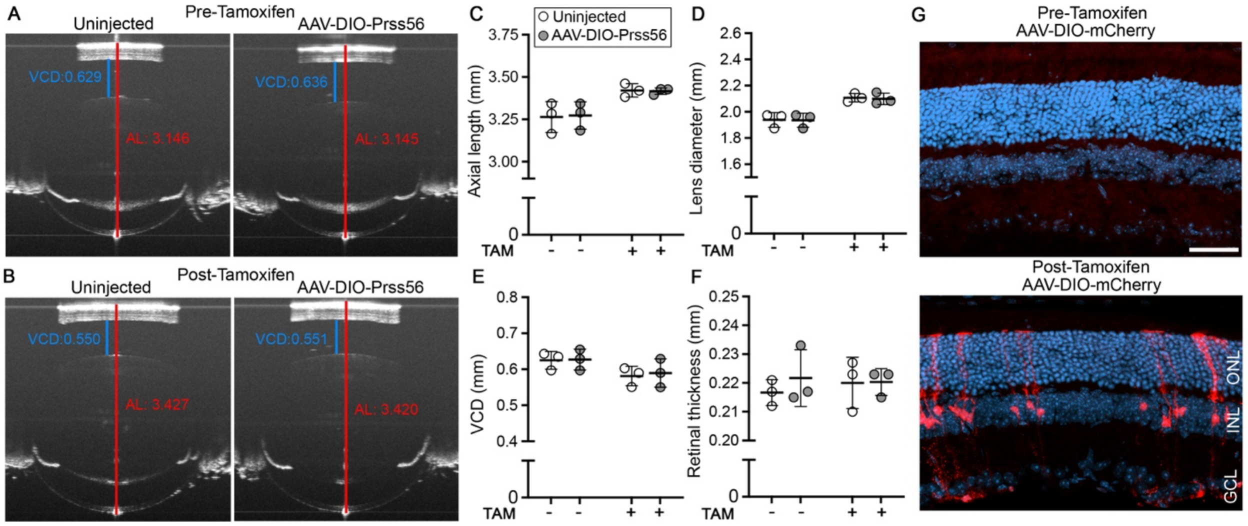


**Supplemental Figure 5. A-F) In absence of Cre, AAV-DIO-Prss56 injection does not alter ocular biometric parameters.** Intraocular injection of AAV-DIO-Prss56 construct in absence of *Glast-Cre*$^{ERT}$ does not alter ocular parameters before or after tamoxifen administration around 1.5 mo. (**A-B**) Representative OCT images and (**C-F**) ocular parameter measurements showing similar axial length and VCD in AAV vector injected eyes and contralateral uninjected control eyes prior to tamoxifen administration (1.5 mo) or in mice aged to 2-3mo following tamoxifen induction. n=3 eyes/group. Data are presented as mean ± SD, paired two-tailed Student t-test. Additional ocular parameter measurements are presented in Supplemental Table 6. **G**) **Representative images from *Glast-Cre*$^{ERT}$ retina showing the presence of tamoxifen-induced mCherry signal in Müller glia in AAV-DIO-mCherry injected eyes.** Scale bar= 50 µm. GCL, ganglionic cell layer; INL, inner nuclear layer; ONL, outer nuclear layer. Additional ocular parameter measurements are presented in Supplemental Table 6.

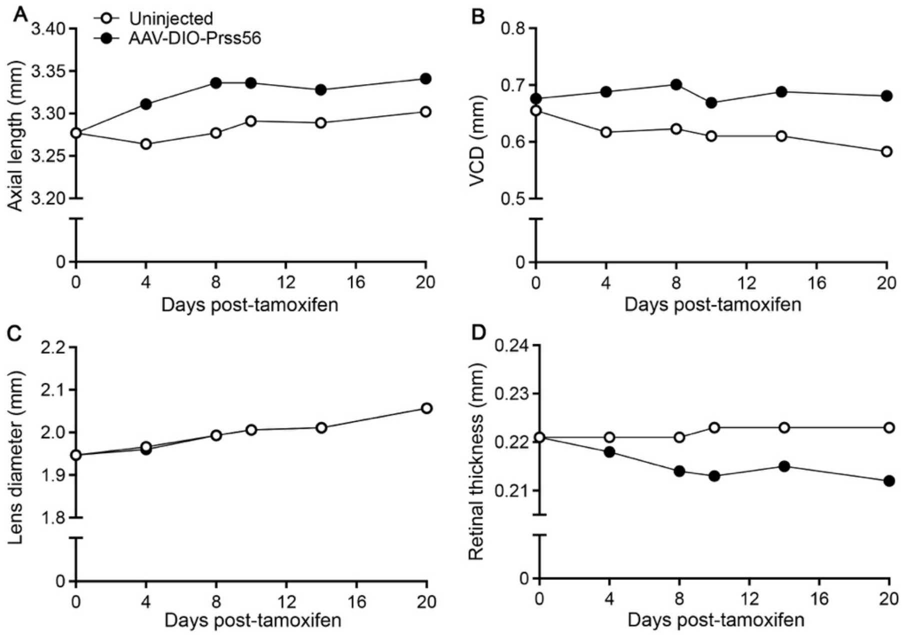


**Supplemental Figure 6. Time kinetics of ocular parameter following AAV-mediated overexpression of *Prss56* in Müller Glia.** (**A-D**) Ocular biometric parameters following tamoxifen administration between AAV-DIO-Prss56 injected and uninjected control at 1.5 mo. Significant increases in ocular axial length (A) and VCD (B) and reduction in retinal thickness (D) were detected in AAV-DIO-Prss56 injected eyes compared to contralateral uninjected control eyes as early as day. Lens thickness remained unchanged between the two groups(C).

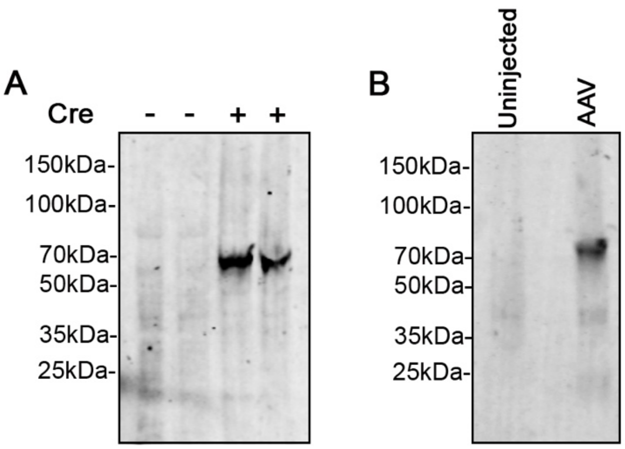


**Supplemental Figure 7. Cre-dependent expression of FLAG-tagged PRSS56 catalytic mutant protein. A)** Western blot confirming expression of the FLAG-tagged PRSS56 catalytic-mutant protein in HEK293 cells transduced with AAV2-DIO-FLAG-PRSS56-Cat Mut in the presence (+) of Cre recombinase. **B)** Western blot showing retinal expression of FLAG-tagged PRSS56 catalytic-mutant protein following intravitreal delivery of AAV2-DIO-FLAG-PRSS56-Cat Mut in Glast-CreERT mice after tamoxifen induction. FLAG was detected using a monoclonal anti-FLAG M2 antibody.

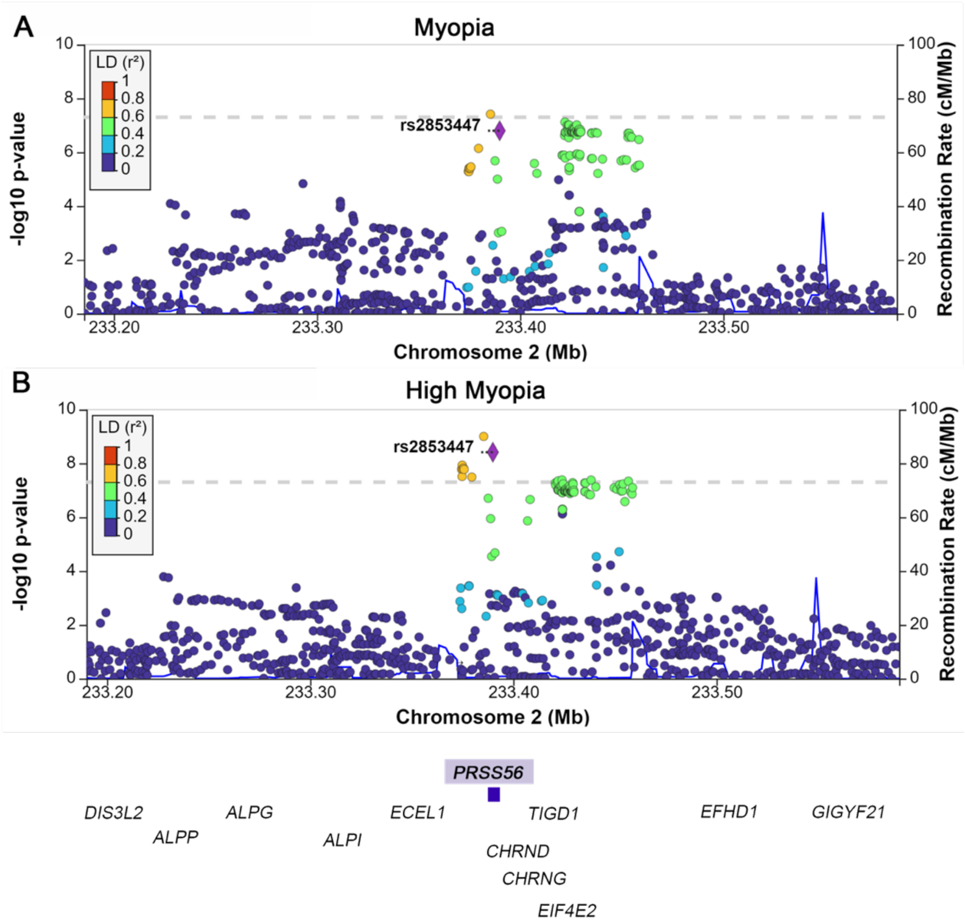


**Supplemental Figure 8.** Locus Zoom plots of 1 MB region around *PRSS56* in GERA non-Hispanic white individuals showing an association of the genetic variant rs2853447 at *PRSS56* locus with all myopia (**A**) and high myopia (**B**).

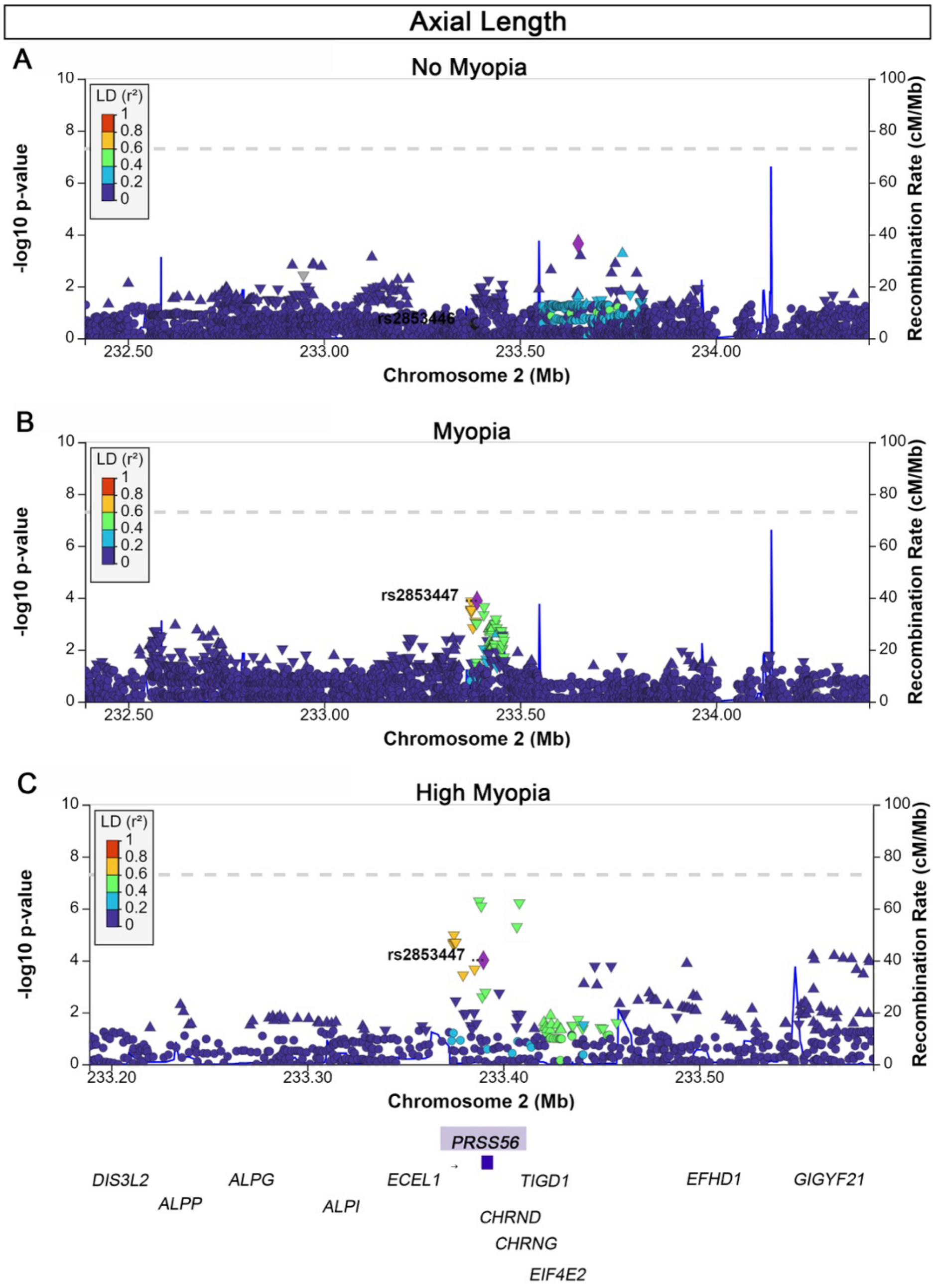


**Supplemental Figure 9.** Locus Zoom plots of the *PRSS56* region showing differential association with axial length in GERA non-Hispanic white participants according to myopia status. (**A**) no myopia (**B**) myopia (all) (**C**) high myopia.

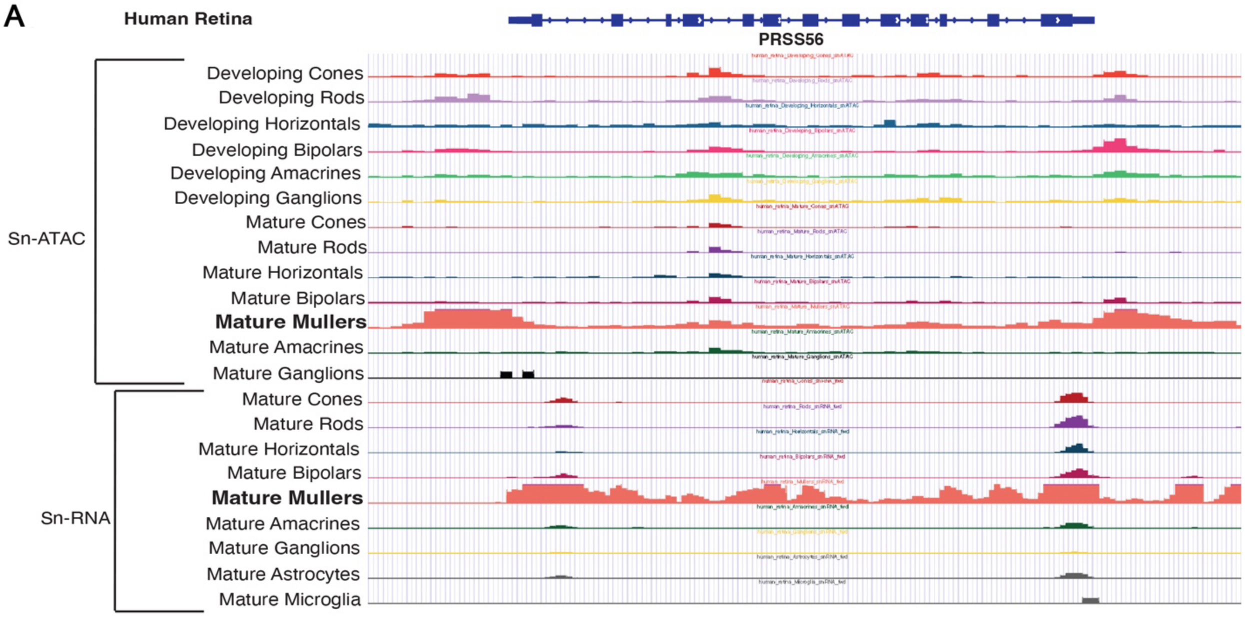


**Supplemental Figure 10**. **Chromatin accessibility and gene expression at the PRSS56 locus across human retinal cell types.** Tracks show single-cell ATAC-seq (snATAC) and single-nucleus RNA-seq (snRNA-seq) profiles from developing and mature human retina. Top panel (Human Retina, sn-ATAC): Cell-type specific chromatin accessibility signals, including progenitors (early, late, cycling), precursors, and differentiated retinal neurons (cones, rods, bipolar cells, horizontal cells, amacrine cells, ganglion cells, and Müller glia). **Bottom panel (sn-RNA-seq):** Cell-type–specific transcriptional activity in mature human 7retinal cell-types reflecting gene expression associated with accessible regulatory regions.

.
